# Accounting for Spatial Correlation in Graphical Analysis of Spatial Transcriptomics Data

**DOI:** 10.1101/2025.07.23.666450

**Authors:** Ana Gabriela Vasconcelos, Patrick Danaher, Daniel McGuire, Jon Wakefield, Ali Shojaie

## Abstract

Co-expression analysis is key for understanding disease mechanisms and gene regulatory and functional relationships. In spatial transcriptomics, estimating gene correlation is challenging due to correlation among cells, which can lead to spurious associations that obscure true biological associations. To address this, we propose SpaceDecorr, a method that adjusts gene expression for technical artifacts and spatial dependencies by modeling each gene independently using a Negative Binomial Generalized Additive Model (NB-GAM) with spatial splines. Co-expression is then estimated from the Pearson residuals, yielding decorrelated expression values suitable for downstream analysis. This method targets cell-intrinsic coordination, rather than clustering genes by shared spatial patterns, and supports multi-sample analysis trough independent per-sample adjustment. Across simulations and real datasets, it consistently reduces false-positive correlations and improves the functional coherence of co-expression modules.

## 1 Introduction

Understanding disease processes and cellular functions requires more than identifying individual marker genes—it demands insight into how genes interact within complex biological networks. These networks, typically modeled with genes as nodes and edges denoting co-expression, regulatory relationships, or shared functional roles, can provide a systems-level understanding of cellular behavior and disease mechanisms [1, 2]. Analyzing gene interactions can reveal disease-relevant pathways, improve disease classification, inform biomarker development, and ultimately support therapeutic discovery [3, 4].

Methods such as Weighted Gene Co-expression Network Analysis (WGCNA) [5] and differential network analysis have been widely used to characterize gene modules and detect context-specific associations in bulk or single-cell RNA-seq data. Gene associations are known to vary across biological conditions—such as treated vs. untreated samples—or across distinct cell types or tissue regions. Examining how these gene associations change in response to experimental, environmental, or pathological perturbations provides valuable insights into cellular functions [6, 7].

Recent advances in Single-cell Spatial Transcriptomics (scST) allow the mapping of gene expression at cellular resolution while preserving tissue architecture. This added spatial context enables not only the characterization of cell-specific gene expression, but also the identification of spatially localized subpopulations, the exploration of microenvironmental differences, and the investigation of cell-cell interactions within native tissue architecture. However, spatial transcriptomics data exhibit multiple sources of variation that can confound inference if not properly accounted for.

Gene expression variation in spatial data arises from two main sources: intrinsic variation and broad tissue-level trends. Intrinsic variation reflects gene expression differences due to cell-internal processes, such as transcriptional regulation, epigenetic state, or cell cycle phase. These internal states often display spatial structure on a smaller scale, with neighboring cells sharing similar gene expression profile if they are in the same cell cycle phase, belong to the same differentiation lineage, or respond to a localized environmental cue. While this variation often reflects relevant biology, it also introduces spatial autocorrelation into the data, which can affect the estimation of gene-gene associations if not properly addressed.

In contrast, broad tissue-level trends, or large-scale variation, arise from expression changes across tissue space. These trends often reflect spatial organization at the anatomical level, such as differences across brain layers or between tumor and stroma, as well as global gradients in signaling or nutrient availability. Large-scale structure can also be driven by technical artifacts, such as batch effects across fields of view (FOVs), increased noise at tissue edges, or gradual shifts in expression due to uneven sequencing depth or tissue handling. Like small-scale variation, these broader trends introduce spatial autocorrelation into the data, but on a larger scale.

These two sources of variation inform different types of biological questions, each requiring specific methodological approaches. The first aims to leverage spatial patterns to identify genes that define tissue architecture. The second seeks to estimate gene-gene associations that reflect intrinsic coordination, such as co-regulation and shared biological functions.

To support the first goal, methods such as SpatialDE [8], MERINGUE [9], COVET [10], SPARK [11], and Giotto [12] aim to uncover spatial co-expression modules by clustering genes with similar spatial patterns—whether small- or large-scale—to identify functionally distinct tissue domains. Importantly, clustering of genes based on spatial patterns may reflect various biological or technical factors: co-expression in the same cell types or regions, a shared response to spatial cues like morphogens or inflammation, participation in the same pathway, coordinated functional activity, or simply technical artifacts like batch effects across slides.

While primarily designed for clustering genes based on spatial patterns, the methods mentioned above are also used to infer gene-gene interaction networks. However, spatial similarity alone does not imply co-regulation or functional interaction, and may instead reflect underlying spatial autocorrelation. Moreover, intrinsic gene-gene interactions may not be discernible from broad tissue-level trends. To bridge this gap, our work focuses on estimation of gene-gene associations that reflect intrinsic coordination in spatial expression patterns.

The key challenge is that spatial autocorrelation can lead to inflated or misleading correlation estimates: when two genes exhibit similar spatial patterns, they might appear correlated even though they are not biologically interacting—a form of spurious correlation. This problem has long been recognized in spatial statistics, where it is known that time and spatial autocorrelation can inflate correlation estimates and lead to false associations. Early work in spatial statistics highlights this problem [13, 14], and subsequent methods have addressed it by removing spatial trends before estimating relationships between variables [15]. More recently, this idea has been extended by considering other data types, such as counts [16], demonstrating how removing spatial structure from data can help isolate true biological effects. Adapting these principles to spatial transcriptomics can improve the accuracy of inferred gene networks by separating true co-regulation from spatial proximity effects.

A few recent methods have been developed to estimate gene-gene correlations in ST data, while accounting for spatial autocorrelation. These include approaches that focus on differences in correlation across tissue regions [17], estimate region-specific or shared co-expression networks [18], or model spatially varying precision matrices [19, 20]. However, these methods primarily focus on within-sample variation between regions and are often tailored to specific use cases.

In practice, many spatial transcriptomics studies involve multiple tissue sections or biological samples, often from distinct individuals or experimental conditions. In these settings, spatial autocorrelation becomes an even greater concern: biological heterogeneity, anatomical variation, and sample-specific technical effects can induce spatial trends that differ across samples. Without appropriately modeling these sample-specific spatial patterns, correlation estimates may be biased, and comparisons across samples may confound biological signal with spatial noise.

To address these challenges, we introduce SpaceDecorr, a method tailored to spatial transcriptomics data that improves the estimation of gene-gene associations by removing confounding spatial structure. The key idea is to adjust for shared spatial trends—particularly broad, tissue-level variation that can drive widespread spurious correlations—while preserving localized, biologically meaningful variation that reflects cell-intrinsic processes and microenvironmental cues. Unlike existing methods, SpaceDecorr supports multi-sample analysis by performing spatial adjustment independently within each sample. This allows for robust and comparable co-expression inference across heterogeneous datasets. Additionally, because it operates as a preprocessing step, SpaceDecorr integrates easily into existing analysis pipelines, enabling researchers to recover more reproducible and biologically relevant gene networks. Through simulations and real-data benchmarks, we show that SpaceDecorr reduces false-positive correlations, improves module interpretability, and enables accurate differential network analysis in spatial transcriptomics data.

## 2 Results

### 2.1 The SpaceDecorr Framework

Here we briefly describe our proposed method. Inspired by existing normalization strategies, we propose adjusting each gene independently for both technical factors (e.g., library size) and spatial effects. This results in a decorrelated dataset, where expression measurements across cells are approximately independent—unlike the original data, which exhibits spatial correlation.

Let *Y_ij_* denote the transcript count for gene *j* in cell *i*, assumed to follow a Negative Binomial distribution with mean *µ_ij_* and gene-specific dispersion parameter *θ_j_* . For each gene *j*, we fit the following regression model:

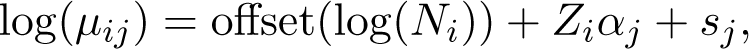

where *N_i_* is the total library size of cell *i*, *Z_i_* is a vector of observed covariates (e.g., batch, quality metrics), and *s_j_* captures the spatial effect for gene *j*. We estimate *s_j_* using a mean-modeling approach via Generalized Additive Models [21] (see Methods for details), although alternative smoothing techniques may also be used.

A key modeling parameter is the basis dimension *K*, which defines the maximum complexity of the spatial function. If *K* is too small, the model may oversmooth the data and fail to capture spatial heterogeneity; if too large, it may increase computational cost without substantial gain. Thus, the choice of *K* represents a trade-off between estimation accuracy and computational efficiency. Results using different values of *K* are presented in Supplementary Figure 1.

After model fitting, we extract the Pearson residuals, which represent normalized gene expression values adjusted for technical and spatial effects. Because each gene is modeled independently, the procedure is naturally parallelizable, allowing for efficient computation in large datasets.

### 2.2 Spatial Structure Inflates Correlations

Spatial transcriptomics data preserve the spatial organization of tissues, but this structure can confound gene-gene association estimates. When two genes are similarly patterned across space— due to anatomical gradients or technical artifacts—they may appear correlated even in the absence of true functional interaction. Such spurious correlations can distort gene networks and mislead biological interpretation. We therefore aimed to evaluate the extent to which spatial structure contributes to inflated co-expression estimates.

We considered the CosMx Non-Small Cell Lung Cancer (NSCLC) dataset [22], which includes 66,267 cells and 960 genes. To focus on reliable signals we filtered out genes with low expression and those prone to segmentation artifacts, yielding a final set of 425 genes. Our final goal is to estimate a gene co-expression networks that reflect functional or regulatory relationships between genes in this tumor subpopulation. To evaluate whether spatial structure induces spurious associations, we constructed a negative control set comprising gene pairs with no known physical or functional interaction (described in STRING or BioGRID) and no shared pathways in curated databases. These gene pairs are unlikely to be biologically connected, so elevated correlations among them would indicate false-positive associations.

Such spurious correlations may arise depending on how gene-gene associations are estimated— particularly in methods that fail to adjust for spatial structure or that incorporate spatial smoothing. Standard approaches from single-cell RNA-seq (scRNA-seq) typically normalize for sequencing depth using log-counts per million (log-CPM) or variance-stabilizing transformations (VST) [23], and then compute correlations on normalized values. While these strategies aim to preserve biologically meaningful variation, they do not account for spatial autocorrelation. Spatial methods such as SPARK and Giotto incorporate spatial information, but are not explicitly designed to isolate gene-gene co-regulation. SPARK applies correlation analysis to VST-normalized counts, similar to scRNA-seq workflows, while Giotto smooths expression across each cell’s *k* nearest neighbors before computing correlations. This smoothing emphasizes spatial structure, both small and large scale, under the assumption that spatial proximity reflects biological coordination.

To illustrate how spurious correlation can arise, consider two genes: *Mitotic Spindle Organizing Protein 2A* (*MZT2A*), a protein coding gene involved in microtubule nucleation and mitotic spindle organization, and *Discoidin Domain Receptor Tyrosine Kinase 1* (*DDR1*), a receptor tyrosine kinase implicated in cell adhesion, extracellular matrix remodeling, and cancer progression. These genes have no known functional or regulatory interactions and do not participate in any common pathways. However, the methods described above estimate a statistically significant correlation between them: log-CPM (*ρ* = 0.115*, p <* 1 × 10*^−^*^20^), SPARK (*ρ* = 0.120*, p <* 1 × 10*^−^*^20^), and Giotto (*ρ* = −0.135*, p <* 1 × 10*^−^*^20^).

Notably, Giotto estimates correlations on spatial effects, and estimated that the two genes vary oppositely to external stimulus, contrary to the others methods. This likely reflects Giotto’s emphasis on spatial structure, as its correlation estimates are based on smoothed expression across neighboring cells. As a result, Giotto’s output is conceptually similar to computing local correlations by aggregating nearby cells: in both cases, most spatial regions exhibit a negative association between MZT2A and DDR1 (Figure 1A). Additionally, visual inspection of the spatial expression patterns of MZT2A and TYK2 (Figure 1B) reveals consistently lower expression of both genes in field of view (FOV) 11, suggesting that shared technical artifacts could also be contributing to inflated correlations across methods.

**Figure 1:**
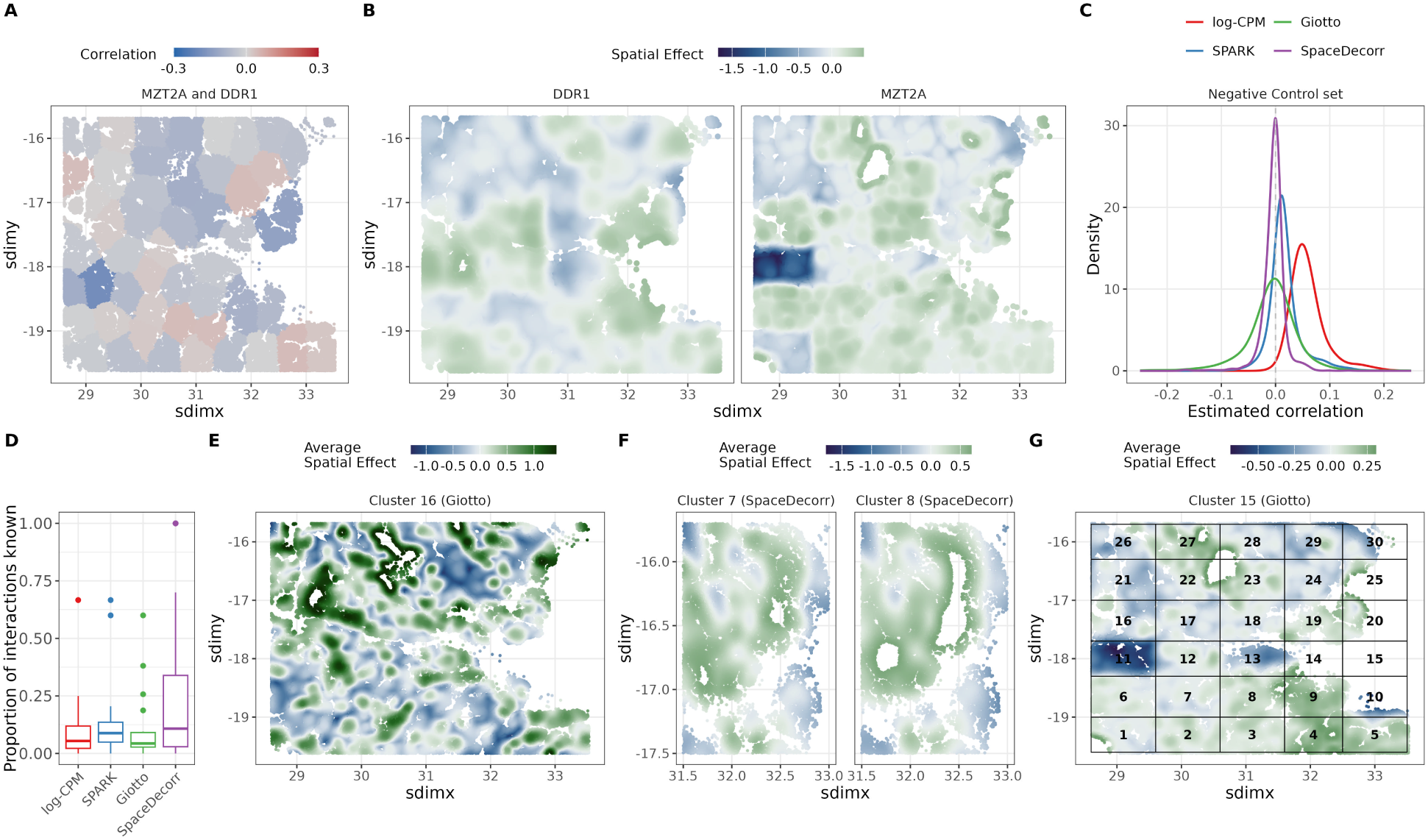
Spatial confounding can inflate gene-gene correlations and obscure biological interpretation. **(A)** Correlation between MZT2A and OSMR computed across spatially grouped cell clusters. **(B)** Spatial expression patterns of MZT2A and OSMR, obtained from the spatial component of a spline-based model. **(C)** Density of estimated gene-gene correlations for pairs with no known protein-protein interaction and no shared pathways, shown for each method. **(D)** Boxplots of biological coherence of modules, described as proportion of interactions known. **(E)** Average Spatial effect for Cluster 16 obtained with Giotto. **(F)** Zoomed in average Spatial effect for Clusters 7 and 8 obtained with SpaceDecorr. **(G)** Average Spatial effect for Cluster 15 obtained with Giotto, overlapped with FOVs.

Applying our proposed method, SpaceDecorr, which decorrelates gene expression while accounting for spatial effects and technical variables (see Methods), we estimate near-zero correlation (*ρ* = −5 × 10*^−^*^5^*, p* = 0.998). This example illustrates how spatial proximity and technical artifacts can confound gene-gene association estimates. Although MZT2A and DDR1 lack any known functional relationship, their co-localized spatial expression leads to inflated correlations in methods that do not explicitly account for spatial dependence.

Importantly, this is not an isolated example. When analyzing all gene pairs with no known protein-protein interactions and no shared pathways, we find that SpaceDecorr report the lowest proportion of significant correlations (39.2%), with estimates centered near zero with the lowest variation (Figure 1C). In contrast, correlations obtained based on log-CPM are almost uniformly positive and shifted toward higher values, suggesting a systematic inflation of co-expression, which resulted in 99.6% of associations significantly different than zero. SPARK reports fewer significant associations than log-CPM (73.5%), likely due to varying sequencing depths, with estimates closer to zero but still shifted towards positive values. Giotto yields correlations centered around zero, but with higher variability, resulting in 76.0% of gene pairs being significant.

These results demonstrate that spatial structure, whether from by biological gradients or technical artifacts can lead to misleading gene-gene associations. While standard approaches such as log-CPM, SPARK, and Giotto partially address technical variability or leverage spatial information, they do not fully correct for spatial dependence. SpaceDecorr more effectively mitigates these confounding effects, providing a robust basis for downstream analyses such as module detection and network construction.

### 2.3 Differences in Co-Expression Modules

We next assessed how differences in correlation estimation affect downstream network analysis. Using the full set of 425 genes, we aim to identify modules of co-expressed genes in tumor cells by applying hierarchical clustering to each method’s correlation matrix. For consistency, we selected 20 modules from each method for comparison (Figure 1E).

To assess the biological relevance of the inferred modules, we examined the proportion of gene pairs within each module that correspond to known physical or functional interactions, based on BioGRID and high-confidence STRINGdb annotations. While median proportions were similar across methods, SpaceDecorr achieved a substantially higher third quartile (0.339) and maximum value (1.0), indicating that its top modules captured more known interactions than those from SPARK (Q3 = 0.136), log-CPM (Q3 = 0.119), or Giotto (Q3 = 0.091) (Figure 1D). In line with these results, pathway enrichment analyses revealed that modules derived from SpaceDecorr tended to recover a greater number and diversity of biologically coherent pathways (352), compared to the other methods (logCPM = 185, SPARK = 210, Giotto = 122).

We proceed on comparing modules obtained with SpaceDecorr, which accounts for spatial dependence, and Giotto, which emphasizes shared spatial trends. While some modules showed substantial overlap, others differed markedly in composition and biological interpretation.

Both Giotto and SpaceDecorr independently recovered a conserved transcriptional module associated with epithelial structure and stress adaptation, corresponding to Cluster 16 and Cluster 13, respectively. This module includes *CRYAB*, *KRT14*, *KRT16*, *KRT6A*, *KRT6B*, and *KRT6C*, and is enriched for biological processes such as intermediate filament cytoskeleton organization, epidermal differentiation, and structural molecule activity. Several of these genes have been implicated in NSCLC progression, including associations with poor prognosis, proliferation, and metastasis [24, 25, 26]. Interestingly, SpaceDecorr also grouped HSPB1 and SFN—assigned to a separate cluster (Cluster 6) in Giotto—into the same module. Both are involved in epithelial stress response and cytoskeletal stabilization; notably, HSPB1 has been implicated in regulating cell survival and therapy response in lung cancer [27]. Spatially, we observed hotspots of higher gene expression localized to specific tumor regions, surrounding areas showing little to no expression (Figure 1E). These low-expression zones may correspond to regions occupied by other cell types, such as immune, stromal, or endothelial cells, which do not express epithelial structural genes. This spatial distribution suggests that the module marks tumor-specific epithelial niches within a heterogeneous microenvironment.

While the epithelial module shows strong agreement across methods, other transcriptional programs are recovered by both methods but with different levels of granularity. For instance, SpaceDecorr identified two transcriptional modules associated with tumor cell proliferation: Cluster 7, enriched for mitotic genes such as *CDKN3*, *MKI67*, and *TOP2A*, and Cluster 8, enriched for DNA replication, repair, and chromatin regulation genes including *BRCA1*, *CHEK1*, *DNMT1*, *PCNA*, and *TYMS*. These clusters reflect distinct stages of the cell cycle—G2/M-phase and Sphase, respectively—and correspond to known proliferative programs in NSCLC [28, 29, 30]. In contrast, Giotto grouped these genes into a single broader module (Module 10), likely reflecting their shared spatial localization and general co-expression in proliferative regions. Spatially, both clusters occupy overlapping tumor regions, but subtle differences—such as regions around sdimx 32–33 and sdimy –17 to –16, where Cluster 7 remains active while Cluster 8 is not (Figure 1F)— highlight specific transcriptional differences that go beyond spatial colocalization. This illustrates how SpaceDecorr can disentangle intra-cellular regulatory signals from spatial proximity, leading to more biologically informative clustering. The comparison highlights how SpaceDecorr provides finer functional resolution, separating biologically distinct transcriptional states that would otherwise be collapsed into a single cluster.

In other cases, Giotto identified modules likely driven by technical artifacts rather than biological signal. For example, Cluster 15, which includes MZT2A and nine other genes, shows no coherent functional enrichment, and notably, MZT2A has no known interactions with any of the other genes in the cluster (Figure 1G). This group displays a distinct pattern of reduced expression specifically in FOV 11, as illustrated previously in Figure 1B. Similar FOV-specific expression differences were also observed in Clusters 2, 4, and 9 (Supp Figure 2). These patterns suggest that such modules reflect technical artifacts, such as batch effects, rather than meaningful transcriptional programs. This example highlights a limitation of Giotto when spatial confounding is not explicitly accounted for.

Together, these comparisons show that both Giotto and SpaceDecorr recover meaningful transcriptional modules from spatial expression data, but they differ in resolution and sensitivity to confounding. Giotto effectively captures broad spatial expression patterns, while SpaceDecorr produces more refined and functionally specific modules, distinguishing programs that may be merged under shared spatial localization. By explicitly removing spatial autocorrelation, SpaceDecorr not only mitigates the influence of technical artifacts—such as field-specific batch effects—but also guards against more subtle, latent spatial biases.

### 2.4 Benchmarking Correlation Estimation Under Simulated Spatial Structure

To assess the accuracy of different methods in recovering gene-gene correlations in spatial transcriptomics data and to validate trends observed in the previous sections, we conducted targeted simulations designed to reflect realistic spatial and expression-level dependencies. We began by evaluating the estimation of pairwise correlations between two genes under controlled conditions.

We simulated multivariate count data using a copula-based framework, ensuring marginal negative binomial distributions while capturing both spatial correlation across cells and correlation between genes (see Methods). To model the underlying expression rate *λ_ij_*, we considered two generative structures: Additive and Matrix Variate (MV).

In the Additive setting (Figure 2A), the expression rate is decomposed into two independent components—one capturing spatial correlation across cells (*Z^s^*) and the other capturing gene-gene correlation (*Z^g^*). This formulation assumes that spatial proximity induces expression similarity independently of gene-gene co-regulation. As such, it isolates the challenge of spurious correlation: genes may appear co-expressed due to shared spatial trends, even if they are not biologically connected. Conversely, true gene-gene correlations are constant across space and not confounded by location.

**Figure 2:**
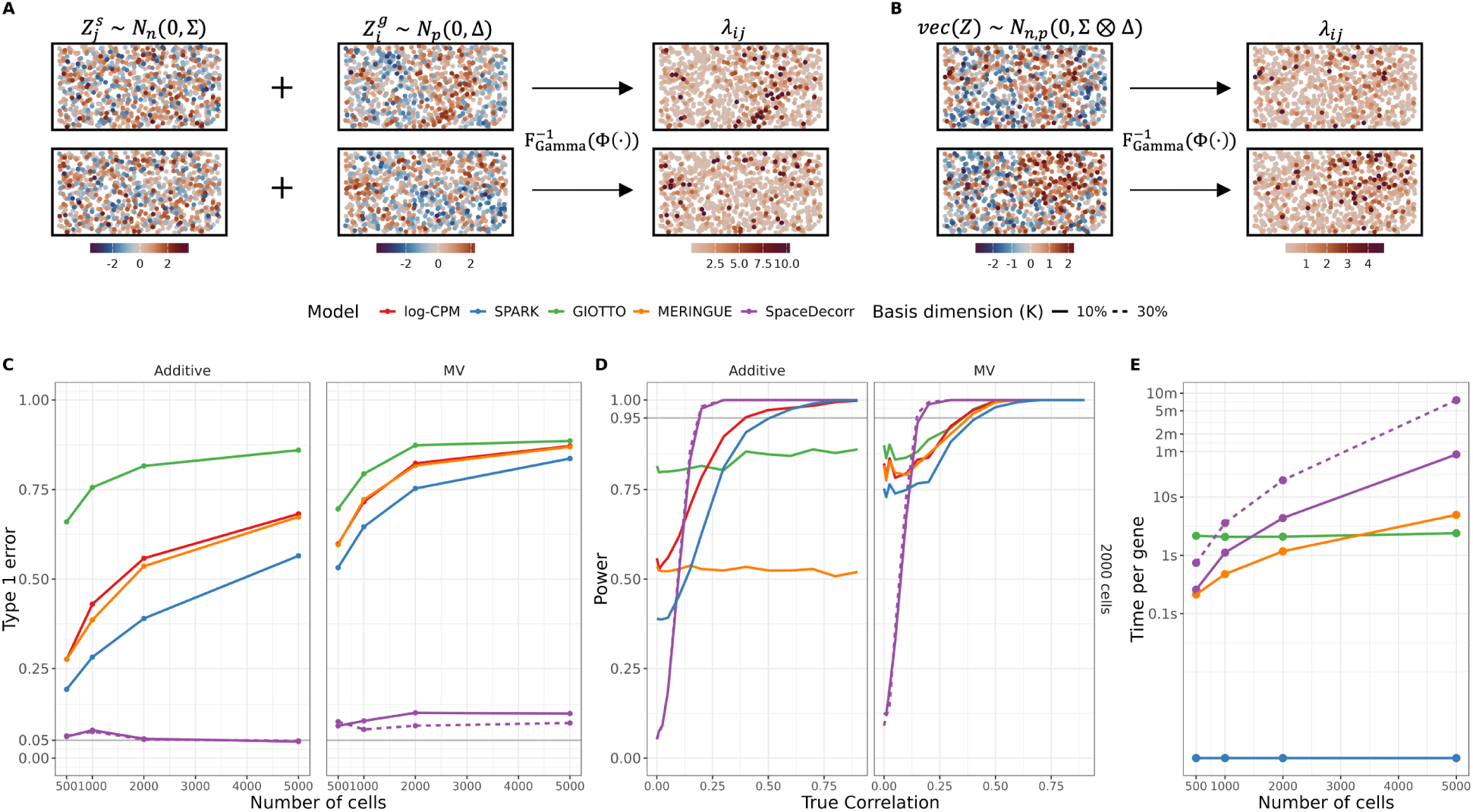
Performance of correlation estimation methods under spatial simulation scenarios. Data was simulated under two frameworks: Additive **(A)** and Matrix Variate **(B)**. Simulation results for testing correlation between two genes across all cells. **(C)** Type I error rates for testing correlation between two genes under the null (true correlation = 0), across increasing numbers of cells (*n* = 100 to 5, 000). Facets correspond to the simulation model: Additive or Matrix Variate (MV). **(D)** Power to detect correlation when the true correlation varies from 0 to 1, under both Additive and MV settings. **(E)** Computational time for each method as a function of sample size.

In contrast, the Matrix Variate setting (Figure 2B) models gene expression using a matrix-normal distribution, where the covariance structure is defined as the Kronecker product of a cell-cell (Σ) and a gene-gene (Δ) covariance matrix. This approach introduces a joint dependency between gene-gene and spatial (cell-cell) correlation. That is, genes that are intrinsically correlated tend to exhibit coordinated spatial expression patterns due to the interaction between spatial and gene-level effects.

Cell coordinates were randomly distributed in a two-dimensional space, and we varied the number of cells across simulation replicates. As the area in which data is simulated remains constant, higher number of cells indicates higher density; that is, more cells in the same area. Spatial correlation was introduced via a distance-based covariance function, and all genes shared the same marginal distribution, with fixed library sizes. After simulating expression data, we compared correlations estimated from SpaceDecorr to those obtained using Pearson correlation on SPARK-normalized and log-CPM-transformed counts. Since library sizes were fixed in this setting, these estimated correspond to correlations computed on the raw and log-transformed data, respectively. We also evaluated correlations produced by existing spatial methods, including MERINGUE and Giotto. For each method, we assessed whether the estimated correlation for a target gene pair was significantly different from zero using Fisher’s z-transformation test.

We also considered a well-established method for gene-gene network estimation, GraphR, which infers partial correlations rather than marginal correlations. Because its objective differs from the pairwise correlation focus of our study, GraphR was not included in the main comparison. Instead, the simulations were adapted to include one independent gene and two correlated genes, enabling a fair comparison between GraphR and SpaceDecorr in a setting suitable for partial correlation estimation. Results are shown in Supplementary Figure 4.

We first evaluated type I error control (Figure 2C). In the additive model, the spatial effects and gene-gene dependencies are simulated independently, allowing us to examine how spatial correlation alone can inflate correlation estimates. We note that even at lower sample sizes SpaceDecorr was the only method to consistently maintain nominal Type I error rates. Other methods that do not directly account for the spatial correlation—SPARK, MERINGUE, log-CPM, and Giotto— exhibited inflated Type I error, with Giotto performing worst, reaching rates above 0.6 across all sample sizes. As the number of cells increased, spatial correlation among neighboring cells also intensified, leading to further inflation of type I error for all methods except SpaceDecorr, which remains robust. Under this simpler model, specifying the spline basis dimension as 10% of the sample size yields performance comparable to higher settings (e.g., 30%), suggesting that moderate basis complexity is sufficient.

These trends persisted under the MV model, which introduces joint dependency between spatial and gene-gene correlation. Here, uncorrelated genes may still exhibit shared spatial patterns due to the coupled structure of the simulation. This stronger spatial structure in this setting led to even greater Type I error inflation for all methods except SpaceDecorr. When using a smaller spline basis (e.g., 10% of the sample size), SpaceDecorr still performed well, though modest gains in type I error control were observed when increasing the basis dimension, emphasizing the importance of flexible spatial modeling under complex conditions. Despite this, SpaceDecorr remained the only method to retain near-nominal Type I error rates control across all sample sizes, with other approaches showing Type I error exceeding 0.25 even at modest sample sizes (e.g., *n* = 500), highlighting their sensitivity to structured spatial dependence.

In addition to controlling Type I error, SpaceDecorr also demonstrated strong statistical power in both the additive and MV settings (Figure 2D). While other methods exhibited inflated Type I error, SpaceDecorr consistently achieved higher power for detecting correlations greater than 0.1. In contrast, methods relying on smoothed counts—such as Giotto and MERINGUE—showed nearly constant power across all correlation values, indicating limited sensitivity and an inability to distinguish weak from strong signals. This suggests that, beyond minimizing false positives, SpaceDecorr improves sensitivity by producing correlation estimates that are less confounded by spatial proximity.

In terms of computational efficiency, all methods completed correlation estimation in under one minute for a sample size of 2,000 cells (Figure 2E). As expected, the runtime for SpaceDecorr increases with larger sample sizes, particularly when using larger spline basis dimensions, which can lead to exponentially longer computation times. However, setting a smaller basis dimension—such as 10% of the sample size—improves computational time by approximately tenfold. For fairness, this benchmark was performed using serial computation across all gene pairs. However, SpaceDecorr supports parallelization, which could substantially reduce runtime and improve scalability for larger datasets.

### 2.5 Alternative Measures of Gene-Gene Correlation

Previously, and throughout this paper unless otherwise specified, we used Pearson correlation to estimate pairwise correlations from the residuals obtained with SpaceDecorr. However, alternative correlation measures may offer advantages in specific scenarios, for example when genes show low expression values. For instance, rank-based Spearman correlation can be more robust to non-linear relationships, while the percentage bend correlation mitigates the influence of outliers by down-weighting the most extreme values (e.g., trimming the top and bottom 10%). These alternatives provide more robust correlation estimates, particularly in the presence of heavy-tailed distributions, outliers, or sparsity, which are common in single-cell data.

To explore the impact of these alternatives, we extended our simulation framework by varying the average gene expression levels—setting the mean to 0.1 to reflect low expression and 0.5 for higher expression—and by including scenarios with negative correlations between genes. These settings help assess how different correlation estimators perform under varying signal strengths and expression sparsity.

For both additive and MV models, when genes exhibited higher average expression levels (mean = 0.5), all correlation methods showed comparable Type I error rates. In these scenarios, both Spearman and percentage bend correlation provided greater power than Pearson for detecting true correlations (Supplementary Figure 5). In contrast, under lower average expression, Spearman correlation resulted in inflated Type I error, while percentage bend correlation maintained error rates similar to Pearson and still achieved higher power. This advantage was especially pronounced when detecting negative correlations.

Furthermore, under the MV model with low expression, estimating negative correlations proved challenging. Negative correlation implies that low expression in one gene is associated with higher expression in the other; however, when both genes tend to be sparsely expressed, this relationship becomes difficult to estimate reliably. These results suggest that filtering out lowly expressed genes may be necessary to ensure stable and accurate correlation estimates.

### 2.6 Module Recovery from Simulated Correlation Networks

After evaluating how each method estimates gene-gene correlation, we next assessed the impact on downstream co-expression analysis. We simulated datasets with 2,000 cells and 100 genes structured into 4 blocks (Figure 3A) representing co-expression clusters. Genes had varying average rate expression, and varying library size, sampled from real data. Correlation was estimated using different methods, followed by hierarchical clustering to recover the underlying clusters. We then compared the inferred and true cluster labels using the Adjusted Rand Index (ARI). Data were generated under both additive and MV frameworks.

**Figure 3:**
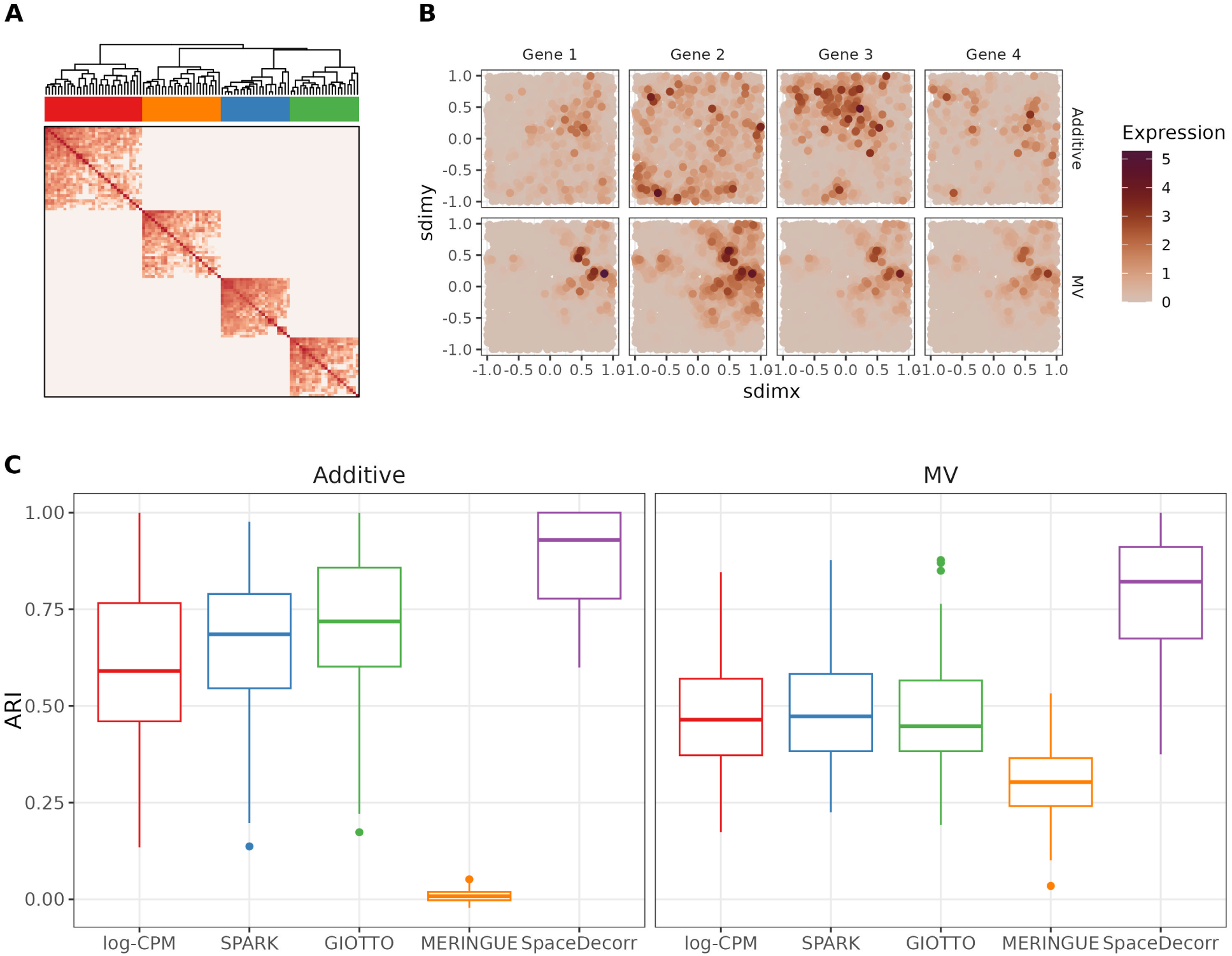
Evaluation of co-expression module recovery using simulated block-structured data. **(A)** Simulated covariance matrix with 100 genes structured into 10 clusters. **(B)** Expression patterns of genes from a single cluster under the Additive and Matrix Variate (MV) simulation settings. **(C)** Adjusted Rand Index (ARI) for cluster recovery across methods, comparing performance with 4 or 10 blocks, with data simulated with Additive or Matrix Variate.

The spatial patterns of gene expression are shaped by the underlying simulation model. Under the Matrix Variate (MV) setting, where spatial (cell-cell) and gene-gene correlations are jointly modeled, genes from the same block tend to exhibit coordinated spatial expression patterns(Figure 3B). This reflects the coupling between correlation and spatial proximity. In contrast, under the Additive setting, spatial correlation across cells and gene-gene correlation are independent. While expression counts still reflect spatial variation due to the shared cell-level component, gene-gene correlation is unrelated to spatial proximity. Consequently, genes within the same cluster may not exhibit consistent spatial patterns.

Under the additive case, where spatial and gene-gene correlations are explicitly modeled as independent components, approaches that rely on spatial co-patterning—such as MERINGUE— performed poorly, failing to recover the true gene modules (Figure 3C). In contrast, SpaceDecorr, which explicitly removes spatial confounding before estimating correlations, achieved the highest ARI, indicating more accurate module recovery due to unconfounded correlation estimates. Giotto and SPARK showed intermediate performance, likely benefiting from partial spatial smoothing but still influenced by residual spatial trends.

In the MV setting, all methods showed lower ARI, as this framework introduces joint spatial and gene-gene correlation, making it more difficult to disentangle spatial effects from genuine coexpression. Nevertheless, SpaceDecorr continued to outperform the other methods, yielding higher cluster recovery. MERINGUE performed better than in the additive setting, as it leverages spatial structure, but still underperformed compared to the other approaches.

Overall, SpaceDecorr consistently outperformed competing methods across both additive and matrix variate models. Giotto and SPARK, despite employing different modeling strategies, showed similar performance, followed by log-CPM and MERINGUE. These results highlight the importance of removing spatial confounding prior to co-expression analysis, especially when genes are more impacted by broad tissue-level variation.

### 2.7 Differential Network Analysis Across Tissue Micro-environments

A key advantage of spatial transcriptomics is the ability to estimate gene networks within distinct tissue niches, offering insight into localized regulatory mechanisms. To evaluate the ability of different methods to recover within-niche networks and detect inter-niche differences, we conducted a simulation study using real spatial coordinates from the CosMx NSCLC dataset. This included 5,878 macrophages spanning two regions: myeloid-enriched stroma (*n* = 4, 594) and stroma (*n* = 1, 281) (Figure 4A). Expression data were simulated under an additive framework for 20 genes, each with varying average expression levels and distinct ranges of spatial correlation decay to better mimic biological variation. Distinct gene-gene correlation structures were assigned across niches, with 25% of the edges modified in one network to induce differential connectivity. An edge in a within-niche graph was defined by a non-zero correlation, while an edge in the differential network reflected a difference in correlation between niches. We investigate two correlation tests, the standard considers Pearson correlation and tests differences with Fisher’s test, the robust considers percentage bend correlation coefficients with a permutation based test. For SpaceDecorr, results using both approaches are presented in the main text, while for other methods only the standard correlation is shown. Full results for all methods using both correlation measures are provided in Supplementary Figure 6. Further details on the simulation setup are provided in the Methods section.

**Figure 4:**
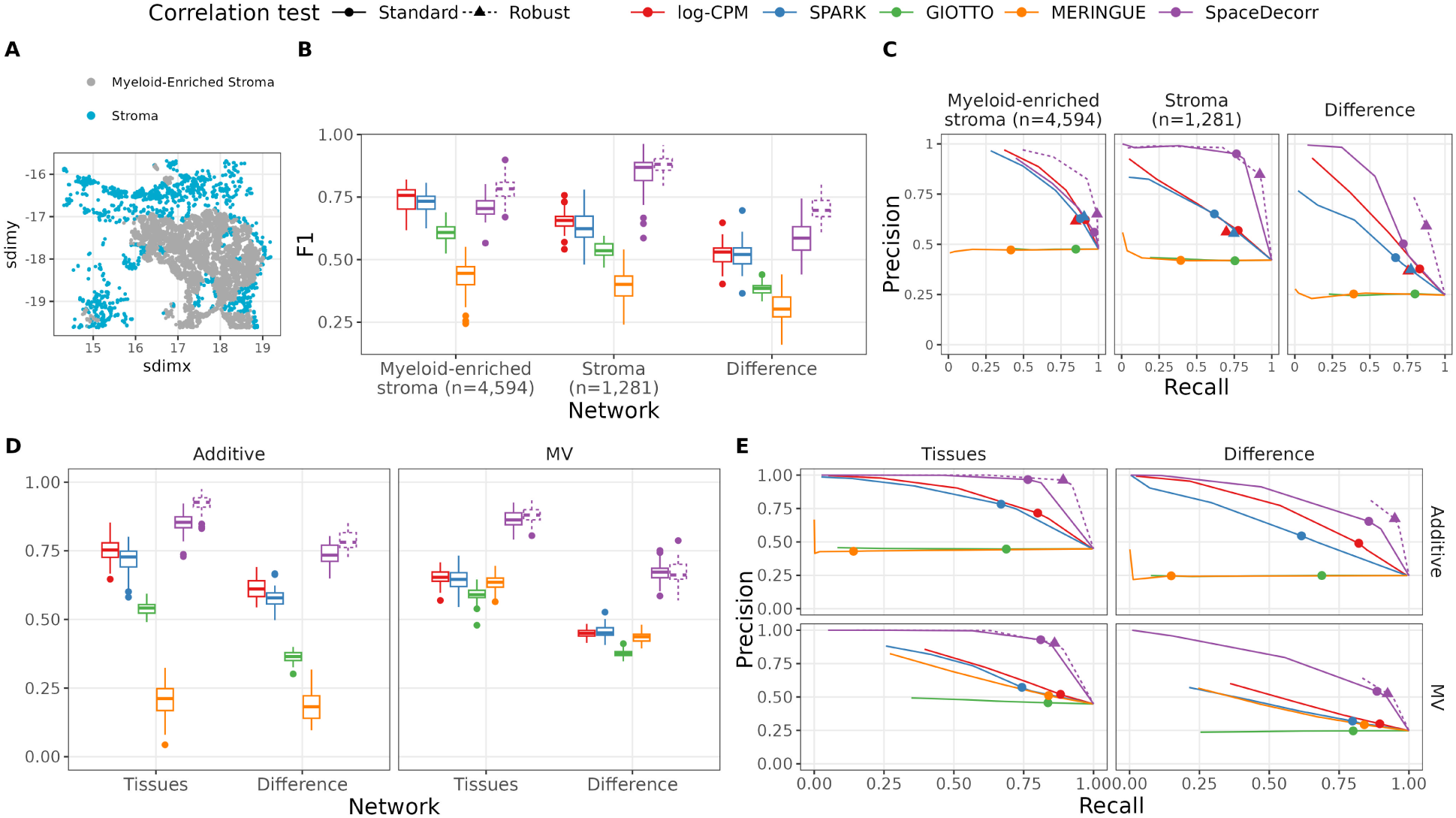
Performance of differential network estimation methods in niche-based and multi-sample spatial simulation scenarios. Simulation results are shown for two settings: (1) network estimation and differential connectivity analysis across two spatially defined tissue niches (**A-B**), and (2) recovery of condition-specific gene networks from spatial transcriptomics data across distinct tissue samples (**C-D**). Line type indicates the test used by SpaceDecorr (Fisher’s Z-test on pearson correlation vs. robust bootstrap correlation test with percent bend correlation). **(A)** Macrophages cells in myeloid-enriched stroma (*n* = 4, 594) and stroma (*n* = 1, 281). **(B)** F1 scores and **(C)** Precision-recall curves for each method in estimating network structure and edge detection within the Myeloid-enriched stroma (n = 4,594), Stroma (n = 1,281), and their differential network. **(D)** F1 scores and **(E)** Precision-recall curves for each method in estimating network structure and detecting differential edges across two spatially distinct tissues, under both Additive and Matrix Variate (MV) simulation models. Points in Precison-Recall curves indicate p-value threshold of 0.05.

We evaluated the F1 score at a p-value threshold of 0.05 as a measure of overall accuracy. In the myeloid-enriched stroma, log-CPM and SPARK achieved average F1 scores of 0.74 and 0.73, respectively, slightly higher than SpaceDecorr using standard Pearson correlation (0.71). Giotto and MERINGUE, which rely on smoothed expression values, performed poorly (F1 = 0.61 and 0.43). When applying robust correlation (percentage bend), SpaceDecorr’s performance improved substantially (F1 = 0.78), while the effect on other methods was not as pronounced (Supplementary Figure 6).

The overall performance declined for log-CPM (F1 = 0.66), SPARK (0.63), Giotto (0.54), and MERINGUE (0.40). In contrast, SpaceDecorr maintained strong performance, with an average F1 score of 0.84 using Pearson correlation and 0.88 using robust correlation—the highest among all methods evaluated. We also assessed recovery of differential network edges. Most methods showed a drop in performance (F1 scores ranging from 0.51 to 0.52), whereas SpaceDecorr outperformed others with F1 scores of 0.59 (standard) and 0.70 (robust).

To better understand these results, we also examined the precision and recall. Giotto and MERINGUE exhibited flat precision-recall curves, indicating limited discriminative power—only a small subset of true edges were recovered, regardless of the p-value threshold used. In the myeloid-enriched stroma, the remaining methods identified a large number of true edges (recall *>* 0.87 at the 0.05 threshold), though with moderate precision (approximately 0.62), reflecting the inclusion of false positives. SpaceDecorr achieved the highest recall (0.97), albeit with slightly lower precision (0.56). Incorporating robust correlation reduced false positives, increasing precision to 0.65 while maintaining high recall (0.99), suggesting improved accuracy in edge recovery.

In the stroma, SPARK showed higher precision than log-CPM (0.65 vs 0.57) but missed more true edges (recall 0.62 vs 0.78). SpaceDecorr achieved the best balance, combining high precision (0.95) and recall (0.77), with recall further improved using robust correlation (precision = 0.85, recall = 0.92).

Differential edge detection proved most challenging. While existing methods achieved high recall (up to 0.85), their low precision (0.34-0.43) led to many false positives. SpaceDecorr again outperformed, with precision and recall of 0.50/0.72 (standard) and 0.59/0.88 (percent bend), making it the only method to maintain both sensitivity and specificity in this setting. Together, these results show that SpaceDecorr, especially with robust correlation, consistently offers the most accurate and reliable recovery of gene-gene relationships across spatial niches and differential contexts.

### 2.8 Condition-Specific Network Inference

To evaluated the ability of different methods to estimate differential gene networks across distinct biological conditions. To this end, we simulated two tissues representing different conditions, each with a unique gene co-expression network. Cell locations were independently simulated for each tissue, with spatial correlations between cells determined by pairwise distances. Expression data were simulated under the multivariate normal (MV) and additive settings. For one tissue, we imposed a small-world graph structure; in the second tissue, 25% of the edges were altered to induce differential connectivity.

Across all settings, SpaceDecorr consistently outperformed other methods. Under the additive simulation model, most methods—including Pearson, logPearson, Spearman, and SPARK—showed comparable performance for within-tissue network recovery (F1 = 0.70-0.78), while Giotto (0.54) and MERINGUE (0.21) performed substantially worse, likely due to their reliance on spatial pattern smoothing. Under the more complex MV model, Giotto and MERINGUE improved (F1 = 0.59 and 0.63, respectively), though Giotto continued to lag behind the other methods. SpaceDecorr achieved the highest F1 scores in both settings for within-tissue graph estimation (0.85-0.93) and was the only method to maintain strong performance in detecting differential edges (F1 = 0.74-0.79 under Additive; 0.67-0.79 under MV). The use of robust correlation further enhanced SpaceDecorr’s accuracy compared to standard correlation, particularly in the additive setting.

From the precision-recall curves, SpaceDecorr showed consistently higher precision within tissues than all other methods, with mean precision of 0.97 (additive) and 0.93 (MV), compared to values ranging from 0.60-0.78 for other approaches. This indicates that SpaceDecorr estimated far fewer false positives while recovering most true edges. When using robust correlation, recall further increased to 0.89 (additive) and 0.86 (MV), without sacrificing precision. In contrast, methods like Giotto, MERINGUE, and logPearson achieved moderate to high recall but at the cost of much lower precision, reflecting a tendency to overpredict edges. For differential edge detection, SpaceDecorr again achieved the best tradeoff (precision = 0.67-0.68, recall = 0.92-0.95 with robust correlation), while most other methods yielded either low precision (less than 0.55) or failed to exceed 0.80 recall.

### 2.9 Lupus Nephritis: Identifying Condition-Specific Co-expression Networks

To investigate disease-associated changes in gene regulation at the cell-type level, we utilized spatial transcriptomics data from Danaher et al. [31], which profiled kidney biopsies from pediatric patients with childhood-onset systemic lupus erythematosus (SLE) using the CosMx Spatial Molecular Imager. The dataset comprises over 400,000 spatially resolved cells from eight SLE patients and four controls. In addition to gene expression, the spatial coordinates, cellular morphology, and cell neighborhood context enabled the precise annotation of 30 distinct cell types using InSituType [32]. Here we focused our analysis on two representative cell types: podocytes, which are essential for maintaining the glomerular filtration barrier, and proximal convoluted tubule (PCT) cells, which are involved in solute reabsorption and dominate the kidney cortex, the region from which these samples were obtained. Podocytes are relatively sparse, with an average of 1,317 cells per tissue (range: 50-6,211), whereas PCT cells are much more abundant, averaging 14,790 per tissue (range: 3,029-46,876). These differences in cell number and spatial organization provide a useful contrast for evaluating the impact of spatial correlation: we expect that spatial confounding will be more pronounced in abundant, spatially structured cell types like PCT.

To identify gene pairs with condition-specific co-expression differences between systemic lupus erythematosus (SLE) and control samples, we performed differential correlation analysis using the DGCA R package [33], applied separately within each cell type.We focused on SPARK and SpaceDecorr for this analysis because both adjust for technical variation across samples and showed superior performance in simulations for recovering biologically relevant correlation differences. Genes were filtered based on mean expression (greater than 0.1 counts per cell), and we excluded those prone to contamination from neighboring cell types. This filtering yielded 162 genes for podocytes and 674 for proximal convoluted tubule (PCT) cells.

To assess the impact of spatial structure on differential correlation results, we compared p-values and correlation estimates obtained from SPARK and SpaceDecorr (Figure 5, Supplementary Figure 7). We also computed average spatial autocorrelation scores for each gene pair (see Methods). In both podocytes and PCT cells, gene pairs with large p-value differences between methods tended to have higher spatial autocorrelation score, while those with consistent p-values had lower spatial structure—supporting the notion that spatial confounding influences co-expression estimates.

**Figure 5:**
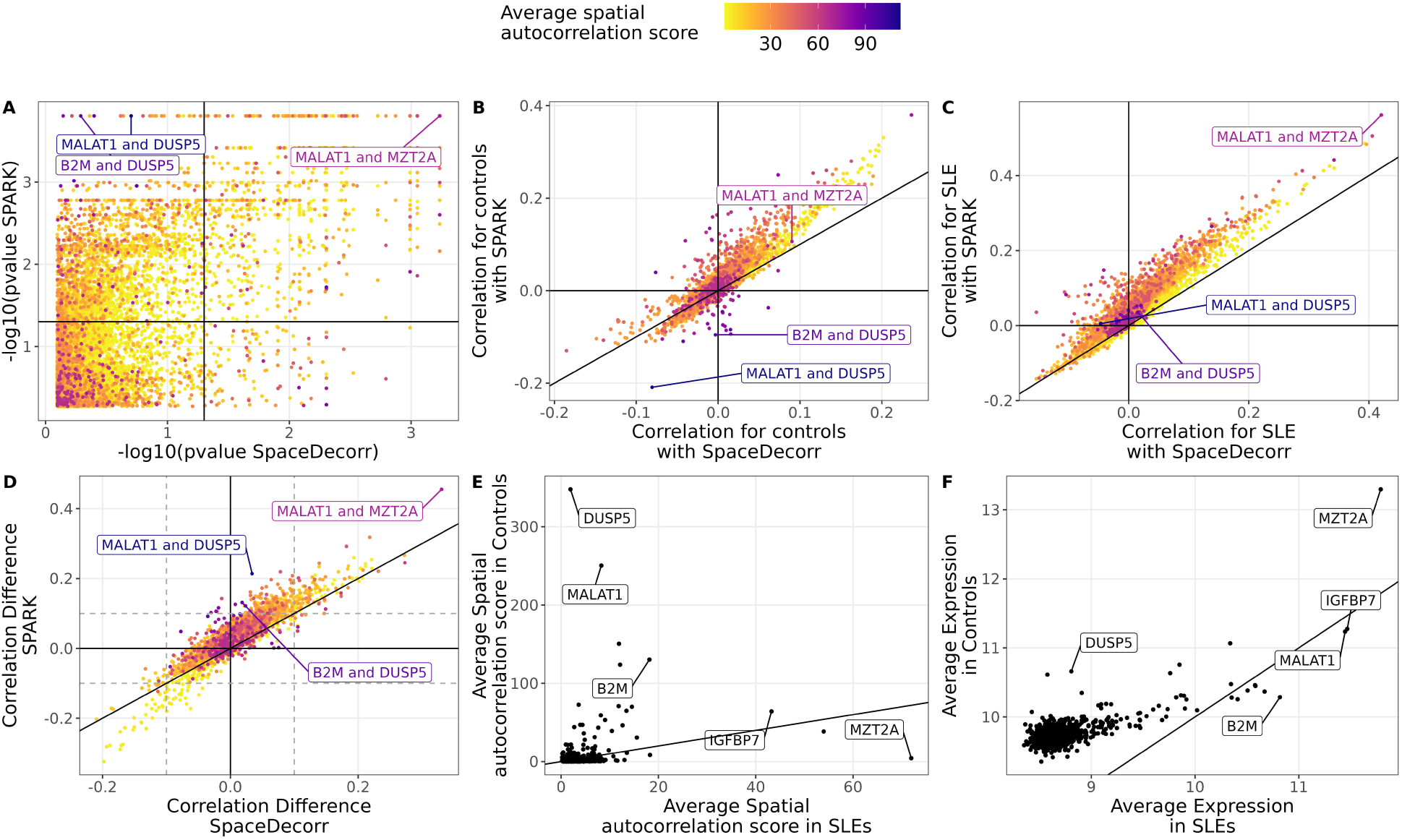
Differential gene correlation analysis between SLE and control samples for Podocytes. **(A)** Comparison of adjusted p-values for gene-gene correlations between SPARK and SpaceDecorr for Podocytes. The solid line denotes the 0.05 significance threshold. **(B-D)** Comparison of gene-gene correlation estimates: **(B)** in control samples, **(C)** in SLE samples, and **(D)** the difference in correlation (SLE - Control). Point color reflects the spatial autocorrelation score, with darker colors indicating stronger spatial structure. **(E-F)** Comparison of gene-level summary statistics between conditions: **(E)** average spatial autocorrelation and **(F)** average expression, each plotted for controls versus SLE.

When comparing correlation estimates directly (Figure 5B–D and Supp Figure 7B–D), we found high concordance between SPARK and SpaceDecorr for gene pairs with low spatial autocorrelation. In contrast, pairs with high spatial structure showed substantial discrepancies: SPARK often over-estimated correlations, while SpaceDecorr returned values closer to zero. Notably, gene pairs with negative differential correlation, more highly expressed in controls, tended to show lower spatial autocorrelation and greater agreement between methods.

Representative examples highlight these patterns. For *B2M* and *DUSP5*, both with high autocorrelation in controls (Figure 5E), SPARK estimated a correlation of -0.09, while SpaceDecorr gave 0.02. A similar case was seen with *DUSP5* and *MALAT1*, with differences in cestimates of correlation between methods. In both, SPARK detected significant differential correlation, but SpaceDecorr did not. In contrast, the pair *MZT2A* and *IGFBP7* —with high spatial autocorrelation in SLE (Figure 5F)—showed divergent correlation estimates (0.08 vs. -0.11), while *MALAT1* and *MZT2A*, both strongly structured, yielded consistent estimates from both methods, showing that for strong signals both methods are consistent.

To investigate condition-specific co-expression patterns in podocytes, we compared differential correlation networks between patients with SLE and controls, focusing on edges with a positive correlation difference greater than 0.1 (gain of correlation in SLE) and statistically significant. The SPARK-based network included 86 genes connected by 398 edges (Figure 6A), while the SpaceDecorr network yielded a more selective set of 47 genes and 132 edges, with 46 genes shared between the two (Figure 6B). Despite this reduction in size, the edge density of the SpaceDecorr network (0.122) was comparable to that of SPARK (0.109), indicating that SpaceDecorr does not simply sparsify the network but instead highlights a more refined subset of interactions, potentially enriched for biologically meaningful changes that remain after adjusting for spatial confounding. We next compared the modular structure and hub gene architecture of each network to evaluate the biological coherence of the retained interactions.

**Figure 6:**
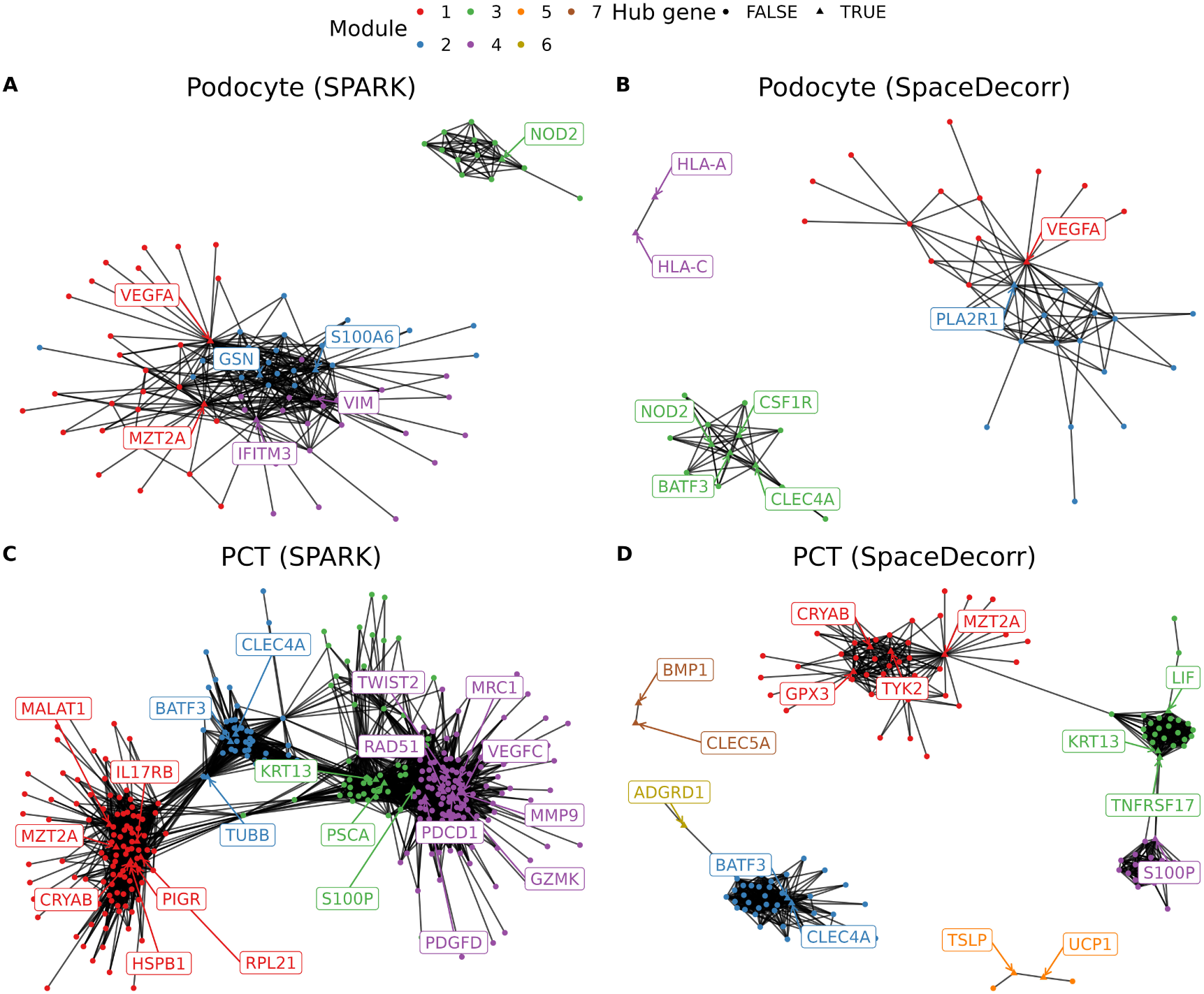
Differential gene co-expression networks in podocytes and proximal convoluted tubule (PCT) cells, estimated using SPARK and SpaceDecorr. Differential co-expression networks were constructed for podocytes (top row) and PCT cells (bottom row) using two spatial methods: SpaceDecorr and SPARK with variance-stabilizing transformation (VST). Edges represent gene pairs showing a gain of correlation larger than 0.1 in systemic lupus erythematosus (SLE) relative to controls.

Both methods identified broadly similar modules, indicating consistent transcriptional responses across approaches. However, network topology differed: SPARK produced a densely interconnected structure with three overlapping modules, while SpaceDecorr yielded a more modular architecture with sharper boundaries. Modules 1 and 3 showed strong agreement in gene composition between methods, supporting the robustness of these SLE-associated co-expression patterns (Supplementary Tables 1 and 2). Module 1 in both networks included a core group of stress-related genes—*VEGFA*,

*DST*, and *CHI3L1* —linked to angiogenesis, cytoskeletal remodeling, and ER stress. These reflect a podocyte-intrinsic stress response commonly observed in lupus nephritis[34]. Module 3, centered on *NOD2*, also overlapped substantially between methods but differed in structure. In SpaceDecorr, it formed a tightly connected immune signaling module, suggesting macrophage activation and immune-podocyte crosstalk, with *CSF1R*, *CLEC4A*, *BATF3* also showing up as hub genes.

A second stress-related module was also shared across methods, centered on *GSN* and *S100A6* in SPARK, and included *CALM2*, *BMP2*, and *HSP90B1*. This module reflects cytoskeletal and calcium signaling changes relevant to podocyte injury [35]. In contrast, SPARK’s module 4, which is deeply connected with modules 1 and 2, grouped interferon response (*IFITM3*), and structural genes (*VIM*), forming a more diffuse module filtered out by SpaceDecorr. Overall, SpaceDecorr retained the core functional modules found by SPARK but with less genes, offering clearer separation of podocyte stress and immune signaling.

To extend our analysis beyond podocytes, we next examined differential correlation networks in proximal convoluted tubule (PCT) cells. Out of the 674 candidate genes, the SPARK-based network showed 337 genes with 4,237 interactions (Figure 6C), while the SpaceDecorr network showed 137 and 918 edges (Figure 6D). As observed in podocytes, the edge density was nearly identical between methods (0.049 for both). Louvain clustering identified four main modules in both networks, enabling direct comparisons of modular architecture and hub gene composition.

Modules 1 and 2 represent largely overlapping biological processes across methods (Supplementary Tables 3 and 4). In Module 1, both methods identify *MZT2A* and *CRYAB* as shared hub genes, implicating cytoskeletal dynamics and chaperone-mediated stress response pathways [36, 37]. SPARK further highlights *MALAT1*, *IL17RB*, *PIGR*, *HSPB1*, and *RPL21* as hubs, pointing to a broader stress-immune response module, with functions involving oxidative stress, heat shock, inflammatory signaling, and epithelial immune defense. In contrast, SpaceDecorr emphasized a more focused subset, including *GPX3* and *TYK2* as additional hubs, shifting the emphasis toward oxidative stress and cytokine signaling [38, 39].

Module 2 features *CLEC4A* and *BATF3* as hub genes in both networks, representing dendritic cell activation and innate immune signaling [40, 41]. SPARK additionally identifies *TUBB* as a hub, a cytoskeletal gene that bridges module 2 with module 1. Many genes in SPARK module 2 form connections with modules 1 and 3, resulting in a less clearly defined module boundary. In contrast, SpaceDecorr’s module 2 appears more topologically isolated, with minimal cross-module edges. This separation may reflect improved discrimination of biological signals, which are often difficult to disentangle from external influences or spatial proximity.

Modules 3 and 4 begin to show stronger divergence between methods. In SPARK, these modules are highly interconnected, with overlapping gene memberships and diffuse boundaries. For instance, *KRT13*, *PSCA*, and *S100P* appear as co-hubs within a shared inflammatory-remodeling module.

SpaceDecorr, however, separates these signals into distinct modules: one centered on *S100P*, indicative of inflammation and calcium signaling, and another incorporating *KRT13*, *LIF*, and *TNFRSF17*, suggestive of adaptive immunity and epithelial signaling[CITE]. These SpaceDecorr modules showed reduced connectivity and clearer boundaries, reflecting a more granular partitioning of biological processes relevant to SLE pathology in PCT cells.

For comparison, applying Giotto to the same data yielded near-fully connected networks: 162 genes with 6,717 edges for podocytes and 674 genes with 169,799 edges for PCT cells (Supp Fig 8). While this approach captures a wide array of interactions, its extreme density results in limited biological interpretability. In contrast, methods that explicitly adjust for spatial structure produce more focused and interpretable networks by reducing spurious associations and emphasizing the strongest, most biologically plausible connections.

In conclusion, SpaceDecorr enables the identification of more distinct modules, each composed of tightly connected genes that are more clearly separated from one another. It effectively filters out genes and connections that may reflect spatial proximity or other extrinsic influences rather than true biological co-expression. This filtering effect is especially pronounced in tissues with high cellular density and stronger spatial correlation such as PCT, but it is still observable in sparser regions, where localized spatial structure can inflate associations like in podocytes.

## 3 Discussion

Spatial transcriptomics (ST) data inherently has two sources of variation: within-cell, gene-intrinsic factors such as regulatory or functional relationships, and broad tissue-level variation that affects groups of genes similarly due to shared spatial or technical influences. While both sources are biologically informative, they serve different analytical goals. In particular, when estimating gene-gene associations, failing to account for spatial autocorrelation can result in inflated or misleading correlation estimates [13, 14]. That is, genes that are not related can appear correlated because of shared spatial patterns.

To address this, we developed SpaceDecorr, a method that removes structured spatial effects via gene-wise GLMMs with spatial splines, providing decorrelated expression values for standard downstream use. Inspired by known prewhitening methods in other spatial data applications [15], and normalizations in scRNA-seq that deal with technical aspects and count data [23], SpaceDecorr is tailored to single-cell spatial transcriptomics data. Because it performs adjustment per sample, SpaceDecorr naturally supports multi-sample ST analyses, ensuring fair and robust comparison across conditions or individuals.

We have demonstrated that spatial autocorrelation leads to spurious correlation both in simulated and real data. In simulations where spatial and gene-gene dependencies were either independent (additive) or coupled (matrix variate), only SpaceDecorr consistently maintained type I error control, even as the number of spatially correlated cells increased. Real data analyses further supported this: in a tumor dataset, SpaceDecorr produced correlation estimates for negative control gene pairs that were centered around zero, in contrast to other methods, which showed widespread inflation. This has implications for downstream analysis, for example, when obtaining correlation modules. SpaceDecorr led to modules with higher biological coherence, supported by known functional and physical interactions, as well as reduced confounding from spatial artifacts like field-of-view, and better recovery—higher ARI—in simulated clusters.

SpaceDecorr also improved differential network recovery—both across niches and in multisample settings. In simulation, it yielded better precision and recall in detecting true changes in correlation structure. But we understand the limitations of testing correlations in spatial transcriptomics with several hundred cells. Nevertheless, in real data, we observed that SpaceDecorr-derived networks were more specific, with fewer genes and interactions, highlighting more reproducible results. SpaceDecorr yielded interpretable clusters—for instance, highlighting stress-adaptive tumor cell states or refined PCT modules in kidney samples with high cellular density—with more clearly separated modules, especially when data were more dense.

One of the key strengths of SpaceDecorr is its flexibility and ease of integration. Because it operates as a preprocessing step, it allows researchers to incorporate it into a wide range of existing transcriptomics analysis pipelines, without bias from spatial correlation or technical artifacts like library sizes. It supports both single-sample and multi-sample analyses, making it applicable to comparative studies involving multiple tissue microenvironments, multiple tissues, individuals, or disease states. By performing spatial adjustment independently for each gene and within each sample, SpaceDecorr enables robust and interpretable comparisons across heterogeneous datasets. The method is also readily parallelizable, facilitating its use in large-scale studies.

At the same time, SpaceDecorr has some limitations. Because it models each gene independently, it may overfit and remove spatial signals that are shared across genes and may be biologically meaningful. Genes with low expression or sparse detection across the tissue are difficult to model, which can lead to unreliable corrections. Additionally, due to count sparsity, the method struggles to estimate negative correlations and performs poorly in estimating partial correlations, which limits its application to conditional dependence frameworks such as precision matrices. The difficulty with negative correlations stems from the fact that when two genes are negatively correlated, one might show lower expression, so sparse expression exacerbates estimation error, making such relationships particularly difficult to recover. Finally, computational cost remains a consideration in large-scale datasets with tens of thousands of genes and cells. However, this burden could be mitigated using faster spatial smoothers or other scalable alternatives.

SpaceDecorr is best suited for single-cell ST platforms, where high spatial resolution creates strong spatial correlations. However, it may also benefit high-plex spot-based platforms with one or a few cells per spot, especially when combined with cell deconvolution techniques. Such information could be incorporated into the model as covariates. Beyond transcriptomics, the approach could be extended to other spatial omics data types, such as spatial proteomics or metabolomics, wherever spatial correlation introduces unwanted bias. Although this study focuses on correlation-based analyses, the SpaceDecorr framework could be adapted for other downstream applications, such as differential expression. This is particularly relevant in multi-sample spatial studies, where spatial heterogeneity across samples can interfere with group-level comparisons. Removing spatial confounding in advance could enable more robust and comparable results across experimental conditions or participant groups.

Importantly, while our goal is to estimate intrinsic gene-gene associations, we recognize that this is only one perspective on spatial transcriptomics. Methods that emphasize shared spatial expression, such as Giotto or other smoothing-based approaches, are well-suited to studying microenvironmental structure, revealing spatially organized processes such as immune cell niches, stromal gradients, or tissue architecture. These approaches are crucial for understanding context-specific gene activity. However, when the objective is to estimate cell-intrinsic co-regulation, or to compare gene networks across conditions or individuals, spatial autocorrelation acts as a confounder that must be corrected. In that context, decorrelation is essential to avoid false-positive associations and ensure robust downstream interpretation. Thus, we view these strategies as complementary— offering distinct but equally valuable insights into ST data.

In summary, SpaceDecorr offers a flexible and generalizable preprocessing solution for spatial transcriptomics data by applying gene-wise spatial smoothing to count-based expression data. By removing spatial and technical confounding, it enables more accurate and biologically meaningful inference of gene-gene relationships. While further development is needed to address sparse gene expression and conditional correlation modeling, SpaceDecorr already improves the reliability and reproducibility of network-based analyses in both simulated and real-world settings.

## 4 Methods

### 4.1 SpaceDecorr Implementation Details

Let *Y_ij_* denote the expression count of the *j*th gene for the *i*th cell. We assume that the counts follows a Negative Binomial distribution, with mean *µ_ij_* and gene specific dispersion parameter *θ_j_* . To normalize the expression data while adjusting for technical covariates and spatial variation, we fit a generalized additive model (GAM) marginally for each gene:

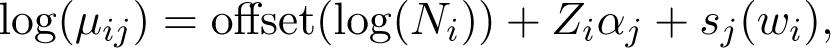

where *N_i_* indicates the total library size for the *i*th cell, *Z_i_* is a vector of observed covariates, such as batch or quality metrics, and *s_j_* (*w_i_*) is a smooth non-parametric function modeling the spatial effect, evaluated at coordinates *w_i_* = (*w*_1*i*_*, w*_2*i*_) ∈ R^2^.

The smooth function *s_j_* (·) is represented as linear combination of basis functions,

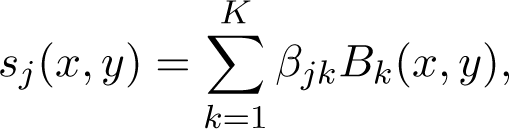

where each basis *B_k_* : R^2^ → R is prespecified, and *β_jk_* is the coefficients associated with the *k*th basis function for gene *j*. Among many possible choices of basis functions, we use thin-plate regression splines [42], though other spline types such as Duchon splines [43] can also be used.

Thin-plate splines estimates *s_j_* (·) by finding the function *f*^^^ that minimizes the penalized loss ∥*y*−*f* ∥^2^ +*λJ* (*f*) where ∥·∥ denotes the usual *ℓ*_2_-norm over the data, *λ >* 0 is a smoothing parameters that controls the trade off between data fitting and smoothness, and *J* (*f*) is a roughness penalty that controls the “wiggliness” of the estimated surface of *f* [21].

To ensure computational scalability in high-dimensional settings, the spline basis is formulated as a low-rank approximation, with the basis dimension *K* chosen to be much smaller than the number of observations, *K* ≪ *n*. This yields an efficient approximation to the smooth function while constraining its maximum complexity. A small value of *K* may lead to underfitting by forcing the function to be too smooth, whereas a large values of *K* increases computational cost without necessarily improving fit. While *K* sets the upper bound on complexity, the actual smoothness is defined by the penalty function *J* (*s_j_*). In our analysis, we choose a value of *K* corresponding to the 10% of the total number of cells to balance computational efficiency and adequate adjustment for spatial correlation (Supplementary Figure 1). However, the optimal choice of *K* may vary depending on the dataset size and spatial resolution.

After fitting the model, the estimated smooth term *s_j_* (*w_i_*) captures the spatial expression pattern of gene *j* at location *w_i_* ∈ R^2^, which can be visualized to reveal spatial structure (as in Figure 1A). To normalize expression while accounting for library size, technical covariates and the spatial effects, we compute Pearson residuals. To avoid the influence of outliers the residuals are clipped at ±^√^*n*, where *n* is the total number of cells [23]. These resulting decorrelated residuals have approximately Gaussian properties and can be used in downstream analysis. Since model fitting is performed independently for each gene, the procedure if naturally parallelized and scalable.

### 4.2 Benchmarking Competing Methods

Throughout the paper, various methods were considered to estimate gene-gene correlation. Log Counts per Million (log-CPM) refers to the Pearson correlation applied to log-normalized counts using the transformation log(10^4^*Y_ij_/N_i_* + 1), where *Y_ij_* is the count for gene *j* in cell *i*, and *N_i_* is the library size for cell *i*.

SPARK [44], originally developed to identify spatially variable genes, was included here by computing correlations on variance-stabilized (VST) [23] normalized counts to allow direct comparison with other methods. We refer to this adaptation simply as SPARK throughout the paper. MERINGUE[45], also designed for identifying spatially variable genes (SVGs), was applied to log-CPM, and spatial cross-correlation matrices were estimated using a spatial neighborhood network with a filtering distance of 0.1. For Giotto [12], the spatial network was constructed using a minimum of *k* = 2 nearest neighbors, and gene-gene correlations were computed using the detectSpatialCorGenes function.

GraphR [19] used VST-normalized data and adjusted for spatial coordinates (*x, y*), estimating correlations at each spatial location. Final correlations were computed as the product of the average estimated correlation and the posterior inclusion probability. P-values were obtained by combining location-specific p-values using the Cauchy combination method.

For niche-specific analyses, spatial methods such as Giotto and MERINGUE were constructed separately for each niche. For GraphR, correlations were estimated at each spatial location and then averaged within each niche. For all other methods, correlations were estimated by restricting the analysis to cells within each niche.

### 4.3 Simulation Setup

We model the expression of gene *j* in cell *i* as *Y_ij_* |*λ_ij_* ∼ Poi(*N_i_λ_ij_*), where *λ_ij_* is the expected expression rate and *N_i_* is the library size for cell *i*. The rate *λ_ij_* is assumed to follow a Gamma distribution, with gene-specific mean *µ_j_* and variance *γ_jj_* . It follows that, marginally, *Y_ij_* follows a Negative Binomial distribution.

Let Σ ∈ R*^n×n^* represent the row (cell-cell) covariance matrix, and Δ ∈ R*^p×p^* represent the column (gene-gene) covariance matrix. The expected rates, *λ_ij_*, are generated using Gaussian copulas in two different frameworks:

- Additive:

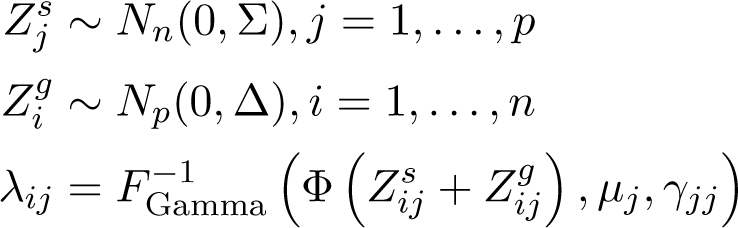

- Matrix Variate:

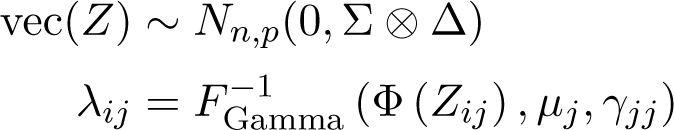

#### 4.3.1 Pairwise Correlation Estimation

To evaluate the estimation of pairwise gene correlations, we simulated spatial coordinates for cells from a uniform distribution in (−1,1) for both *x* and *y* locations. Cell-cell covariance matrix Σ was set to a Matèrn matrix with range 0.5 and smoothness 0.5. We simulated datasets with varying sizes of 500, 1,000, 2,000, 5,000 and 10,000 cells. For each simulation, we generated two genes with correlation varying from 0 to 1. For the comparison with GraphR, three genes were generated: one independent and two others with correlation varying from 0 to 1. Both additive and matrix-variate frameworks were considered, with all genes having Gamma parameters *µ_j_* = 0.5*, γ_jj_* = 1, and fixed library size *N_i_* = 200. To test whether the correlation between two genes was significantly different from zero, we used the test statistic 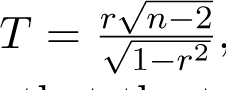, which follows a t-distribution with *n* − 2 degrees of freedom under the null hypothesis that the true correlation equals is zero, where *r* denotes the sample correlation between two genes.

#### 4.3.2 Gene Modules from Correlation Networks

To evaluate the module recovery using co-expression networks, we conducted another simulation designed for WGCNA. Cell locations were simulated uniformly in the (−1, 1)^2^ square. Expression counts for 100 genes were generated, grouped into 5 or 10 network modules. The gene-gene covariance matrix Δ was simulated from using a degree-corrected stochastic block model, similarly to data generation in [46]. We assumed an average node degree of 20, out-in ratio of 1/1000, and a within-community low-degree proportion of 0.9. Node degree followed a power-law distribution with exponent 5. Correlations corresponding to the edges were generated from a Uniform distribution between 0.5 to 0.9, and the resulting matrix was converted to the nearest positive-definite matrix while preserving sparsity using the algorithm from [47].

To simulate expression heterogeneity, the mean expression levels *µ_j_* for each gene were sampled from 0.25, 0.5 and 1, and gene-specific variance was determined using mean-variance ratios sampled from {0.25, 0.5, 0.75, 0.90}, capturing a range of realistic expression dynamics. library sizes were sampled from Lung 5 sample in the NSCLC CosMx dataset. From the simulated expression data, we computed gene-gene correlation matrices and applied hierarchical clustering to recover module structure. Recovery accuracy was assessed using the adjusted rand index (ARI) by comparing the inferred modules to the ground truth.

#### 4.3.3 Niche-Specific Differential Network Analysis

This simulation study aimed to assess whether correlation networks can be accurately estimated for distinct regions of tissue. We focused on the additive model due to the lack of a straightforward way to extend the matrix-variate framework to settings with group-specific covariance structures. Cell locations were based on real dataset of 5,878 macrophages, including 4,597 cells from the myeloid-enriched stroma and 1,281 in the stroma. A shared spatial component *Z^s^* was used for all cells, with the shared Matèrn covariance matrix Σ. In addition, for each gene we generated niche-specific *Z^g^* components, with group-specific gene-gene covariance Δ*_k_* .

To simulate expression data, we modeled 20 genes. For each gene, we sampled a spatial range parameter from the set {0.5, 1, 1.5, 2}, to reflect the diversity of spatial decay rates observed in practice. Mean expression rate were sampled from {0.15, 0.25, 0.5, 1}, and variance was derived from mean-variance ratios sampled from {0.1, 0.25, 0.5, 0.75}. Library sizes were taken as the observed value for each location.

Underlying gene-gene networks were constructed separately for the two regions. In the myeloid-enriched stroma region, used a Watts-Strogatz small-world model with expected neighborhood of 4 and rewiring probability 0.1. For each edge, partial correlations were sampled uniformly between 0.5 and 0.7. To ensure positive definiteness of the precision matrix, we subtracted the smallest eigenvalue from the diagonal. For the stroma region, we changed 25% randomly selected edges by either deleting an existing edge (setting correlation to zero) or creating and edge with correlation sampled randomly, simulating differential network structure between niches.

To estimate genetic networks, we first computed correlation matrices separately for each niche using the normalization methods described previously. Within each niche, we tested whether each pairwise correlation was significantly different from zero using either Fisher’s correlation test or percent bend correlations, in which 10% of extreme values are down weighted to improve robustness [48]. To identify differences in gene-gene associations between niches, we applied the Fisher z-transformation to test for equality of correlations. In addition, to obtain more robust estimates of differential correlation, we used the twocor function from the WRS2 package, which applies a bootstrap-based test for differences between ercent bend correlations correlations. P-values for all tests were adjusted for multiple comparisons by controlling the false discovery rate (FDR), with adjustments performed separately for each niche and for the differential test. The total number of comparisons was based on the number of unique gene pairs. Gene pairs with FDR-adjusted p-values below 0.05 were considered significant and included as edges in the estimated network.

Finally, we evaluated graph recovery by comparing estimated graphs for each region, and on the network difference, against the ground truth. Metrics such as F1, precision and recall were computed to assess accuracy in detecting region-specific networks and their differences

#### 4.3.4 Multi-Sample Differential Network Analysis

This simulation aimed to assess differential correlation network analysis between two distinct tissues or samples. For each tissue, 2000 spatial coordinates were independently generated from a uniform distribution in (−1, 1) for both *x* and *y* axes, following the same setup as the pairwise correlation simulations. Library sizes were sampled separately for each tissue based on real data distributions. Gene expression counts were simulated independently for each tissue using both the additive and matrix-variate copula methods. For the underlying gene-gene correlation structure, we used the same strategy as in the niche-specific differential network analysis: one tissue’s network was generated using a Watts-Strogatz small-world model with an expected neighborhood size of 4 and rewiring probability of 0.1, and in the second tissue, 25% of the edges were randomly modified by either removing existing edges or adding new ones with randomly sampled correlations.

To estimate and compare networks, we first normalized gene expression using either log normalization, the variance-stabilizing transformation (VST), on our proposed SpaceDecorr method, applied separately to each tissue. Gene networks were then estimated following the same procedure used in the niche-specific differential network analysis, including correlation matrix estimation, hypothesis testing for edge inclusion, and FDR correction for multiple comparisons.

### 4.4 Real data

#### 4.4.1 NSCLC tumor dataset

We analyzed the publicly available CosMx Spatial Molecular Imager (SMI) NSCLC FFPE dataset developed by NanoString Technologies. This dataset includes eight formalin-fixed paraffin-embedded (FFPE) lung tumor sections from five non-small cell lung cancer (NSCLC) patients, imaged with subcellular resolution using a 960-gene targeted RNA panel [22]. For our analysis, we focused on Lung6, one of the eight available tissues. We selected tumor cells across multiple spatially defined microenvironments, including tumor-stroma boundaries, tumor interior, tumor cells in stroma, myeloid-enriched stroma, plasma-blast enriched stroma, and general stroma, totaling 66,267 tumor cells.

To reduce contamination-related artifacts and enhance biological interpretability, we filtered the original 960 genes using the overlap ratio metric from the smiDE package [49]. Genes with an overlap ratio greater than 1 (indicative of potential spatial contamination) were excluded. We further removed genes with low expression, defined as having an average expression less than 0.1, yielding a final gene set of 425 genes. We inferred gene co-expression networks using correlation matrices generated from four approaches: SpaceDecorr (*k* = 10%), Giotto, SPARK, and log-transformed CPM (logCPM).

To evaluate the biological coherence of the inferred gene relationships, we compiled reference information from multiple external databases: known gene interactions from BIOGRID [50], interaction scores from STRINGdb (version 12) [51], and pathway annotations from MSigDB [52]. A negative set of interactions was defined as gene pairs that shared no annotated pathways, had no recorded BIOGRID interaction, and had a STRINGdb score of zero or were not found in the database. In contrast, high-confidence interactions were defined as those meeting all three of the following: a BIOGRID interaction, greater than 25% pathway overlap, and a STRINGdb interaction score greater than 700.

Gene modules were identified by applying hierarchical clustering to the correlation matrices, considering 20 clusters. For each module, we computed: the average STRINGdb interaction score across all gene pairs, the proportion of gene pairs with BIOGRID interactions, the average number of shared pathways per pair. In addition, we performed pathway enrichment analysis using the clusterProfiler package [53], evaluating enrichment across all Gene Ontology categories. To characterize the spatial activity of each module, we also calculated the average expression of all genes in the module per cell.

#### 4.4.2 SLE kidney dataset

We utilized publicly available multi-sample spatial transcriptomics data from Danaher et al. [31]. This dataset comprises CosMx Spatial Molecular Imager (SMI) profiles of kidney biopsies from eight pediatric patients with childhood-onset systemic lupus erythematosus (SLE) and four controls. The dataset present expression measurements for 957 transcripts across more than 400,000 spatially resolved cells. For this analysis, we restricted our focus to two epithelial cell types of interest: proximal convoluted tubule (PCT) cells and podocytes.

To improve signal quality and reduce spatial contamination, we applied filtering criteria based on both expression and contamination metrics. Specifically, we retained genes with average expression greater than 0.1 and an average contamination ratio (from the smiDE package) less than 1. This resulted in 674 genes for PCT and 162 genes for podocytes.

To quantify spatial localization patterns, we calculated spatial autocorrelation scores using the binSpect function from the Giotto toolbox. This method identifies genes whose high expression is enriched in spatially adjacent cells and assigns a score indicating the strength of spatial clustering. A score of zero indicates no spatial structure, while positive values indicate spatial coherence. We computed scores for each gene across tissues and cell types, then averaged them across tissues. For visualization of gene pairs, we summarized by averaging scores from both genes in each pair.

Gene–gene correlation estimates unconfounded by spatial structure were obtained with SpaceDecorr. For comparison, we also evaluated correlation matrices based on Giotto’s spatial smoothing and SPARK’s variance-stabilized (VST) normalized data. Correlations for each method were calculated using the DGCA package, using Pearson correlation coefficients and p-values computed separately for SLE and control samples. False discovery rate (FDR) correction was applied, and gene pairs with adjusted p-values below 0.05 were considered significantly correlated. Differential correlation networks were constructed by identifying gene pairs that were differently correlated and showed a correlation increase of more than 10% in SLEs compared to controls.

## Supplementary Material

### Figures

**Supp Fig 1:**
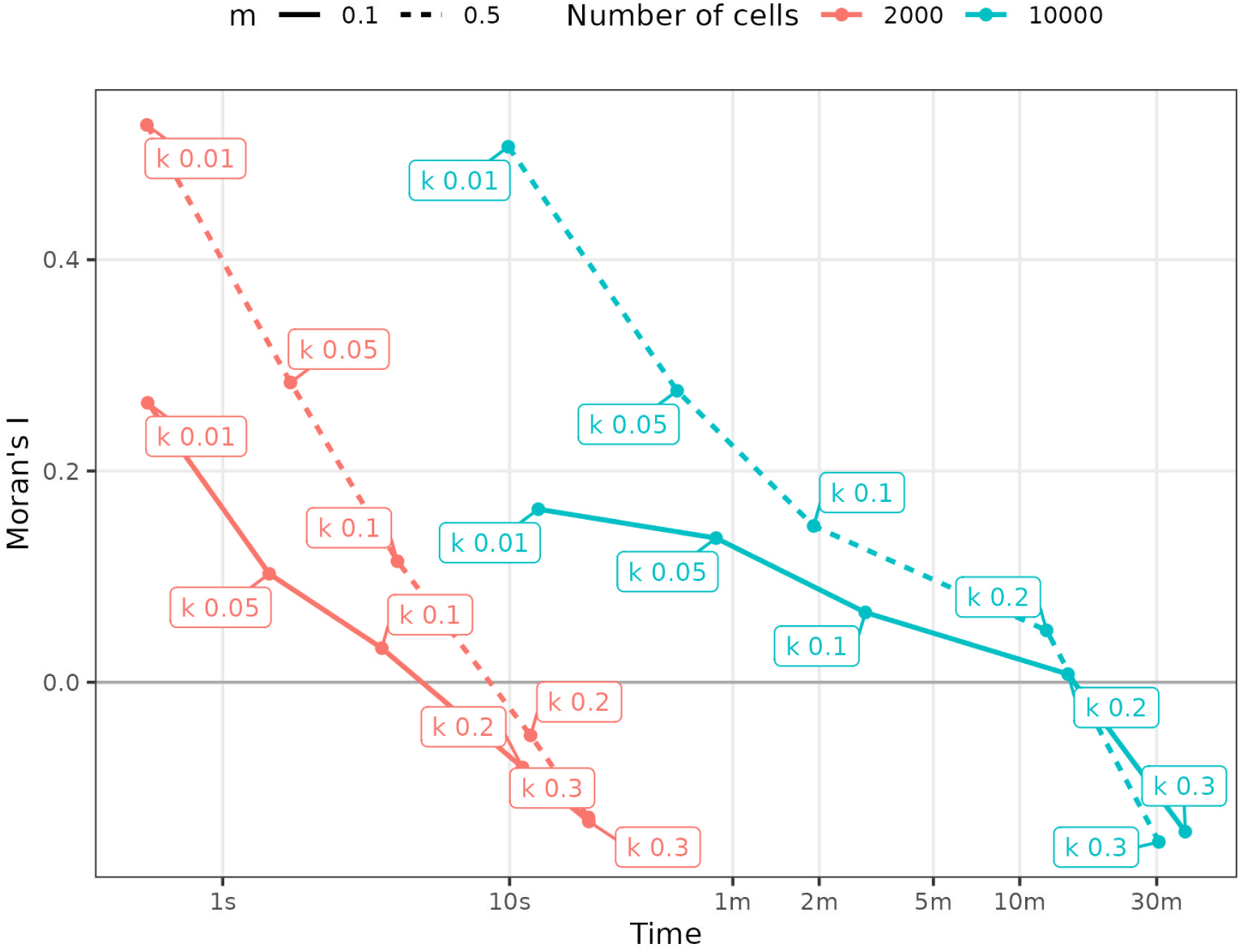
Effect of spline basis dimension on residual spatial autocorrelation and computation time. Considering data simulated under the MV framework, we assessed performance of the model across varying basis dimensions, number of cells and expression levels. Basis dimensions were set to 1%, 5%, 10% and 20% of the total number of cells (i.e., *k* = 0.01, 0.05, 0.1, 0.2). For each configuration we compared the average processing time, and Moran’s I of the residuals, which quantifies unaccounted spatial autocorrelation (values ¿ 0 indicate remaining spatial structure, while values near 0 suggest successful removal of spatial effects). Models were fit under two scenarios of average gene expression: lox (*m* = 0.1) and high (*m* = 0.5), across increasing cell counts. Results indicate that higher basis dimensions are necessary to adequately capture spatial patterns when expression levels or cell density are high.

**Supp Fig 2:**
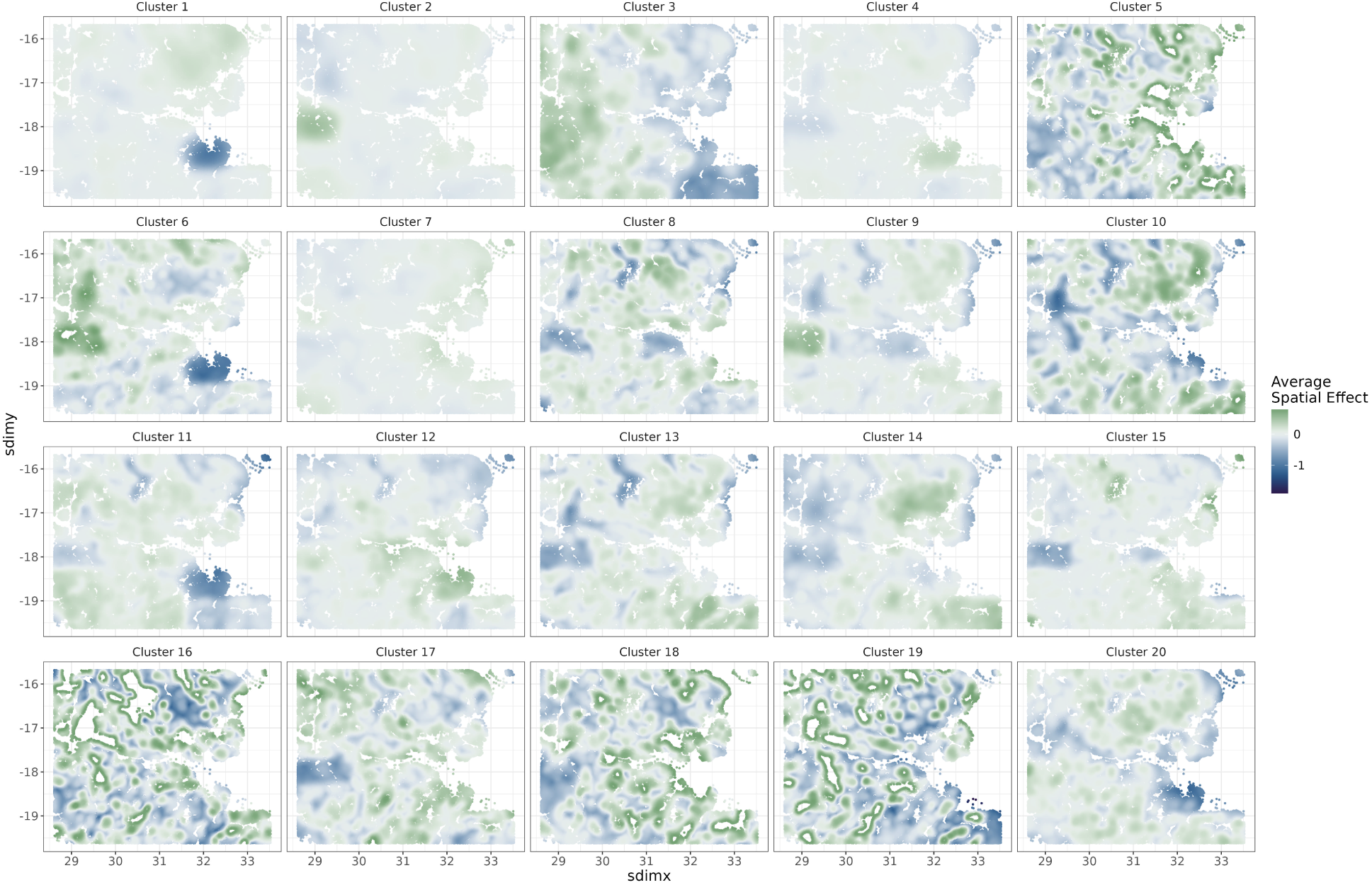
Average spatial expression of all clusters obtained with Giotto.

**Supp Fig 3:**
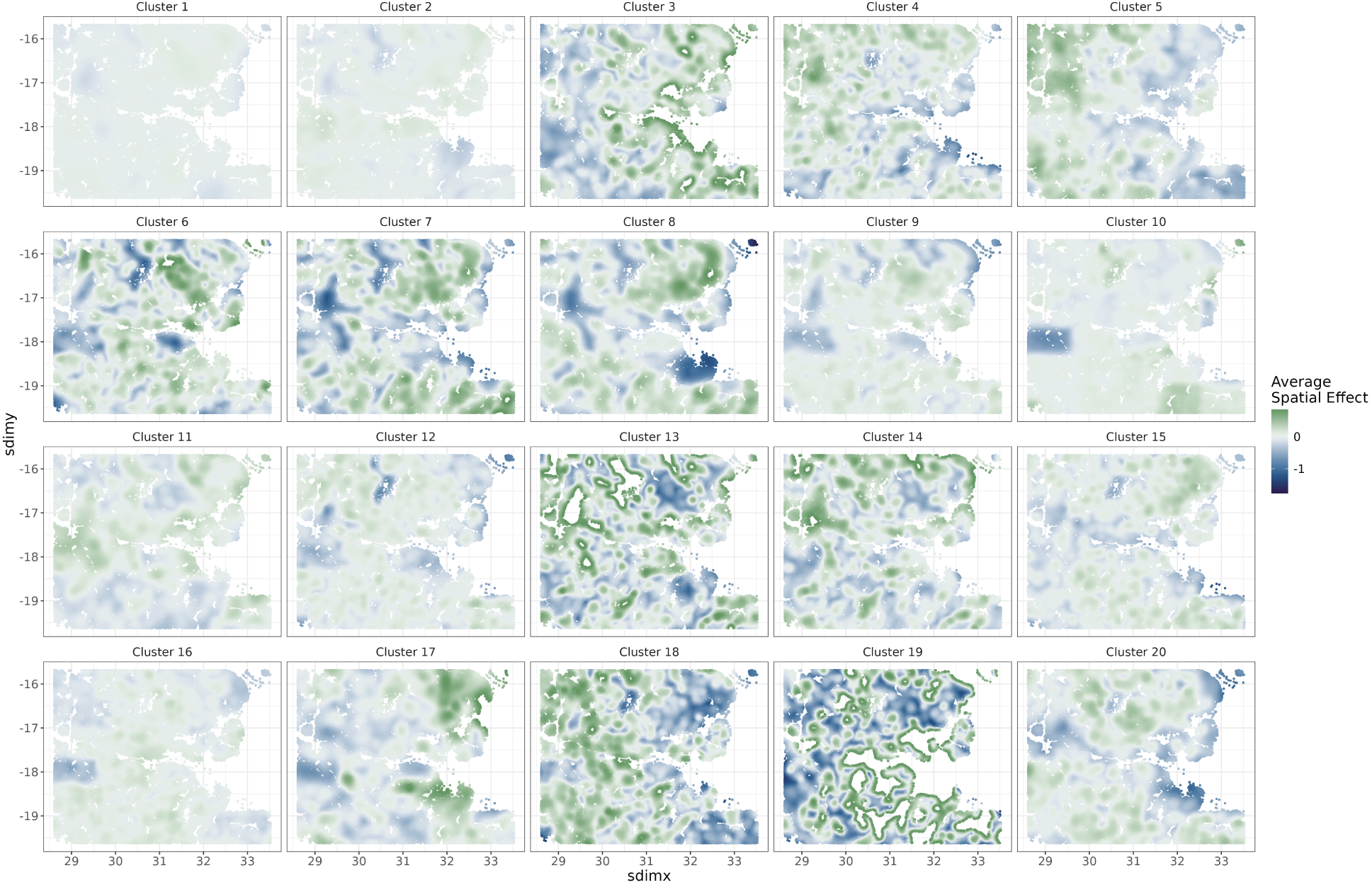
Average spatial expression of all clusters obtained with SpaceDecorr.

**Supp Fig 4:**
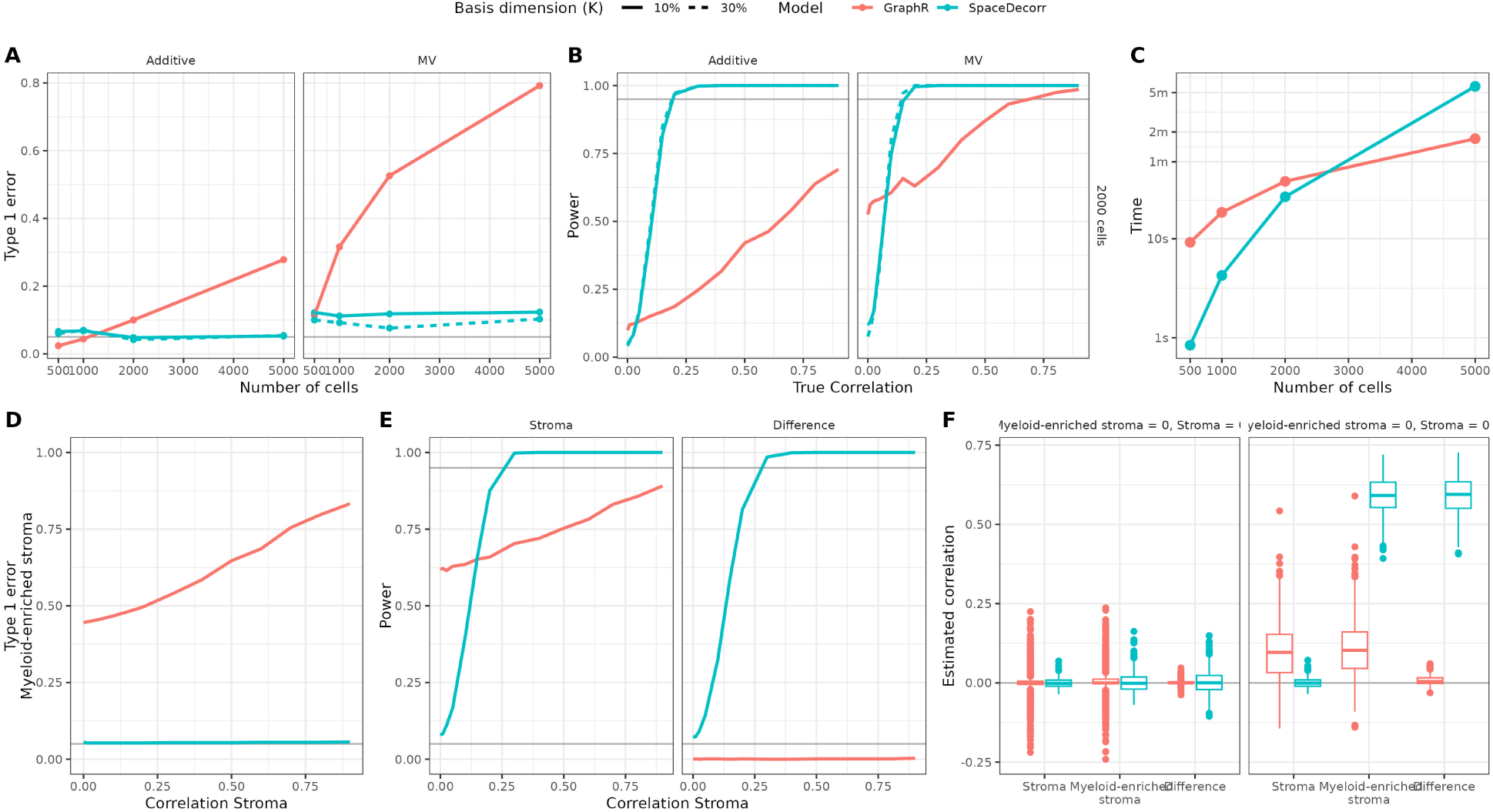
Performance of correlation estimation methods under spatial simulation scenarios with GraphR. Simulation results are shown for two settings: (1) correlation between two genes across all cells (**A-C**), and (2) differential correlation between two tissue niches with distinct correlation structures (**D-F**). Each panel summarizes results across 1,000 simulation replicates. **(A)** Type I error rates for testing correlation between two genes under the null (true correlation = 0), across increasing numbers of cells (*n* = 100 to 5, 000). Facets correspond to the simulation model: Additive or Matrix Variate (MV). **(B)** Power to detect correlation when the true correlation varies from 0 to 1, under both Additive and MV settings. **(C)** Computational time for each method as a function of sample size. **(D)** Type I error for testing correlation in the Myeloid-enriched niche (where the true correlation is 0), across varying correlation values in the Stroma niche. **(E)** Power to detect correlation in the Stroma niche (left) and power to detect a difference in correlation between niches (right), as the true correlation in the Stroma increases. **(F)**Estimated gene-gene correlations are shown for each niche and their difference across methods, under two settings: (1) both niches have true correlation of 0 (null case), and (2) the myeloid-enriched stroma has true correlation 0 and the stroma niche has true correlation 0.9.

**Supp Fig 5:**
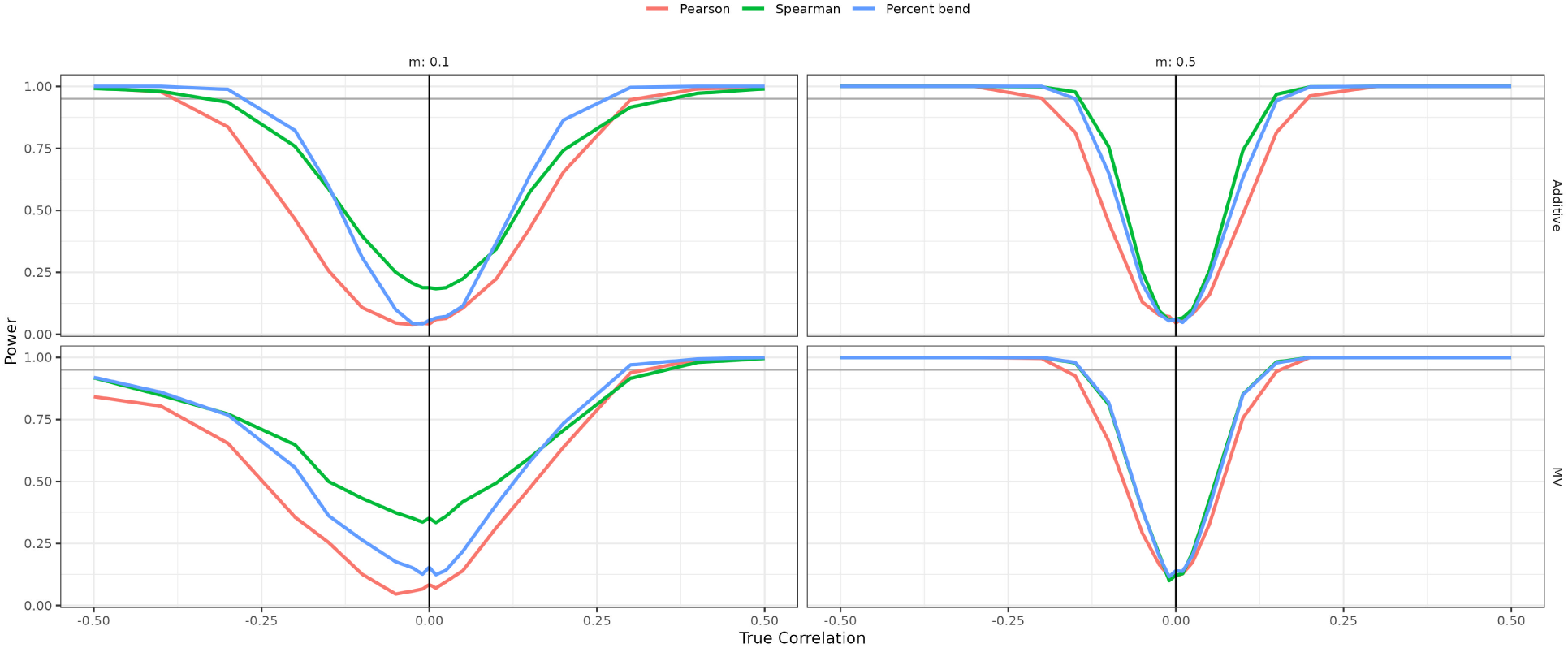
Comparison of correlation metrics for gene-gene association testing using SpaceDecorr. We evaluated the power to detect true gene-gene correlations using Pearson, Spearman, and percentage bend correlation coefficients computed on residuals from SpaceDecorr. Simulations were conducted under two spatial models (Additive and Matrix Variate) and two average gene expression levels (*m* = 0.1 for low expression, *m* = 0.5 for higher expression). For each setting, we varied the true underlying correlation from 0 to 1 (or -1 to 0 for negative correlations), and estimated the proportion of simulations (out of 1,000 replicates) in which a significant correlation was detected at *α* = 0.05.

**Supp Fig 6:**
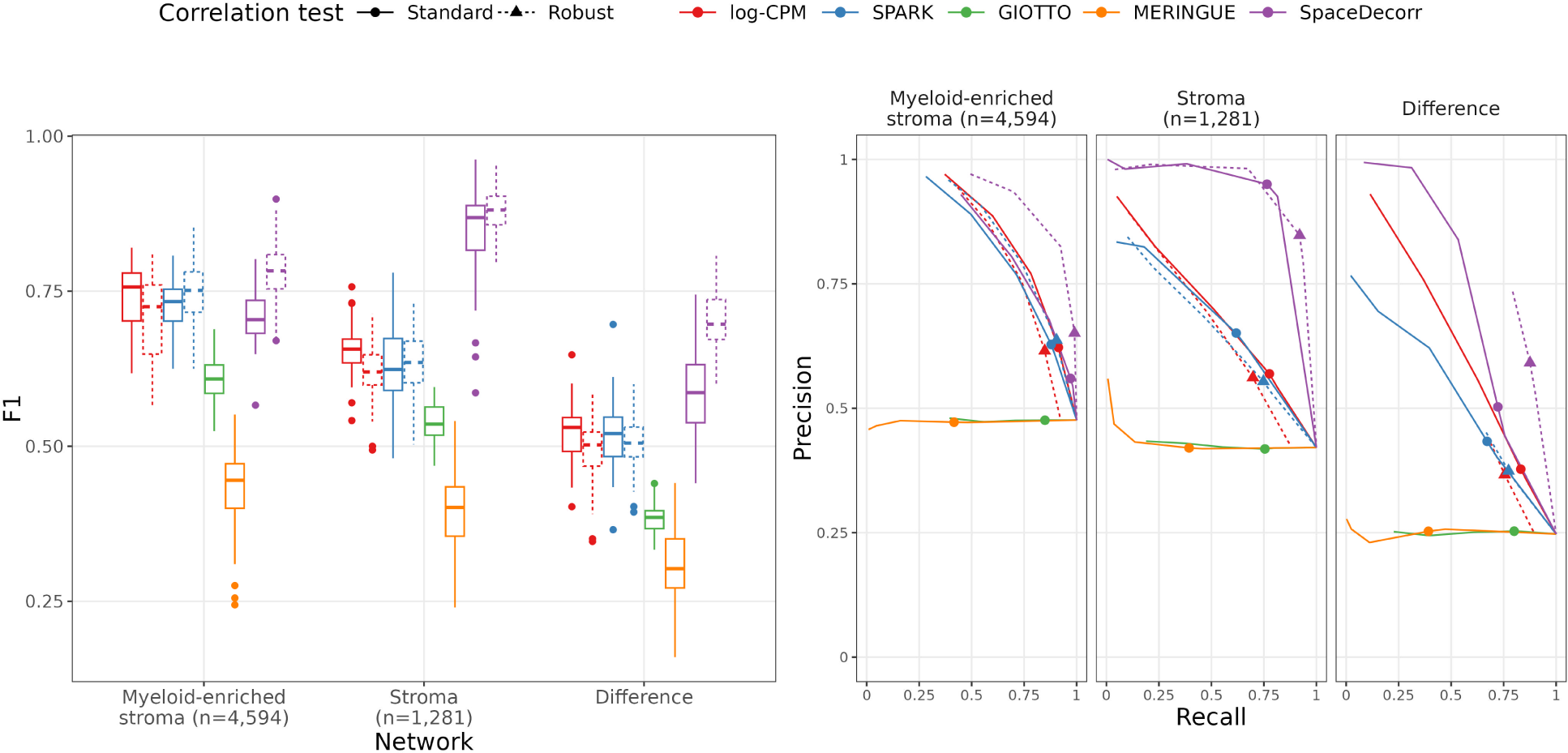
Performance of differential network estimation methods in niche-based spatial simulation scenarios. Simulation results are shown for network estimation and differential connectivity analysis across two spatially defined tissue niches. Line type indicates the test used by SpaceDecorr (Fisher’s Z-test on Pearson correlation vs. robust bootstrap correlation test with percent bend correlation). **(A)** F1 scores and **(B)** Precision-recall curves for each method in estimating network structure and edge detection within the Myeloid-enriched stroma (*n* = 4,594), Stroma (*n* = 1,281), and their differential network.

**Supp Fig 7:**
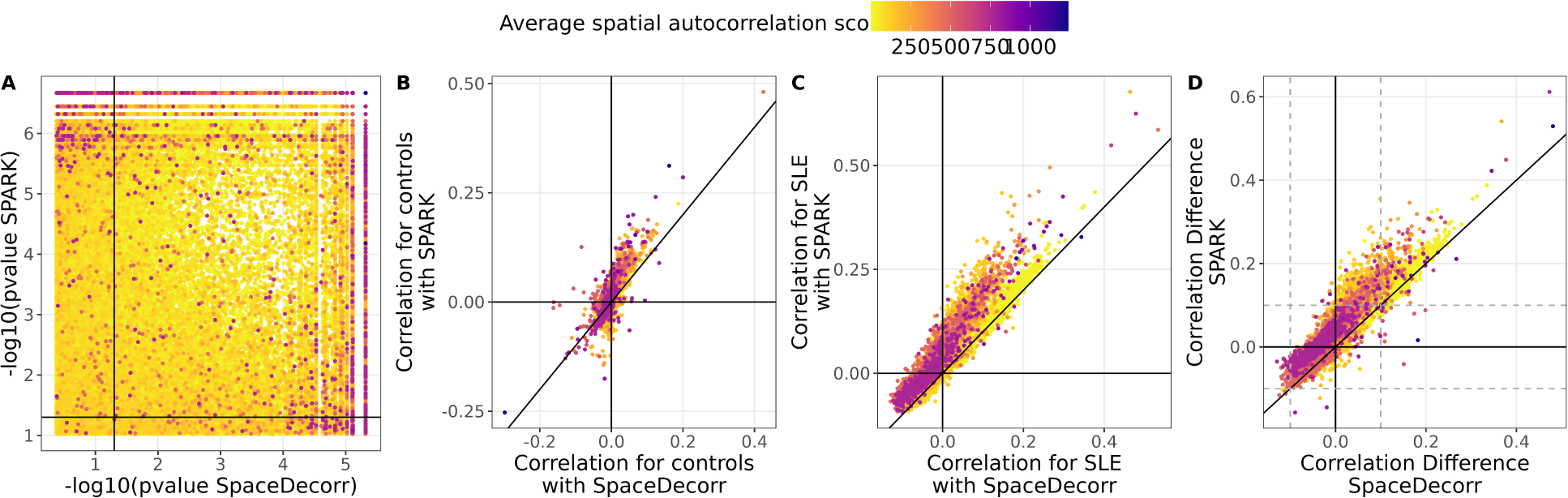
Comparison of gene-gene correlation estimates and significance across normalization methods in PCT. (A) Comparison of adjusted p-values between SPARK and SpaceDecorr. The solid line denotes the 0.05 significance threshold. (B) Comparison of gene-gene correlation estimates in control samples. (C) Comparison of gene-gene correlation estimates in SLE samples. (D) Comparison of correlation differences between SLE and controls. The solid diagonal line indicates zero difference; dashed lines mark ±0.1. In all panels, point color represents the spatial autocorrelation score, with higher values indicating stronger spatial structure. Positive values in (D) reflect gain of correlation in SLE, and negative values reflect loss of correlation.

**Supp Fig 8:**
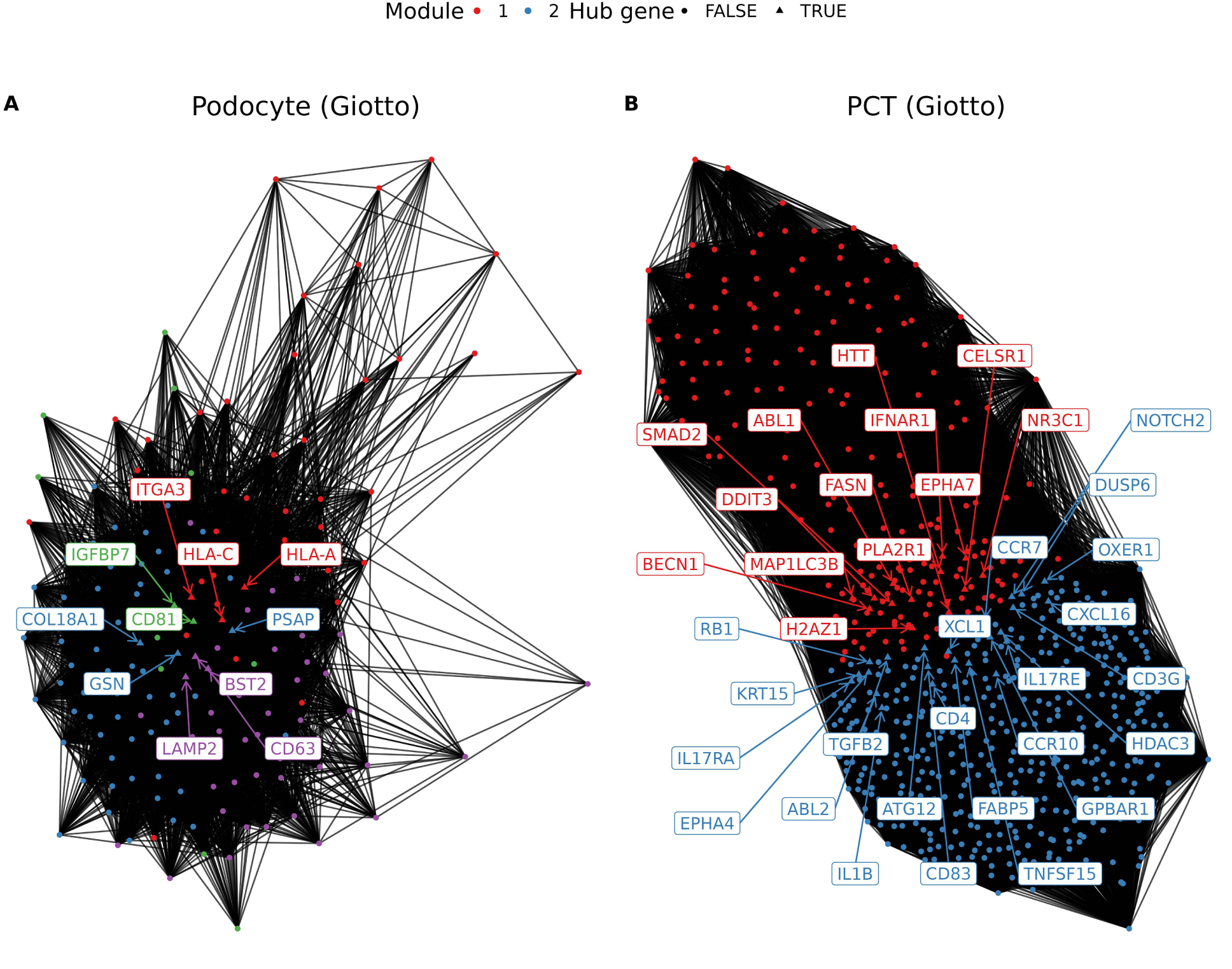
Differential gene co-expression networks in podocytes and proximal convoluted tubule (PCT) cells, estimated using Giotto.

### Tables

**Supp Table 1:** Hub genes and corresponding modules identified using Louvain clustering on the SPARK-inferred gene network in podocytes

| Module | Hubs | Genes |
| --- | --- | --- |
| 1 | VEGFA, MZT2A | MALAT1, MZT2A, DUSP5, CD3E, DST, DCN, PTGDS, VEGFA, MYL9, CHI3L1, COL4A5, MTOR, EPHA2, HSPA1A, TNFSF14, NPR3, HILPDA, DUSP1, FKBP11, RARRES2, HSPA1B, EZR, FYN, FGFR1, PTGES3, TYK2, TPM2, SMO |
| 2 | S100A6, GSN | PLA2R1, GSN, NEAT1, ITGA3, ANXA2, ITGB8, S100A6, ITGAV, SPOCK2, CD81, MXRA8, CALM2, ST6GALNAC3, HSP90B1, APOD, BST2, ITGB5, NDRG1, EFNB1, S100A4, RAMP3, LGALS3BP, MRC2 |
| 3 | NOD2 | CSF1R, BATF3, CLEC4A, NOD2, CD28, NPR2, CLOCK, TUBB, TLR2, NR1H2, SMARCB1, EPHB6, YBX3 |
| 4 | IFITM3, VIM | LGALS1, IGFBP7, HLA-A, B2M, IFITM3, HLA-C, S100A10, BMP2, PSAP, CD63, COL18A1, TIMP1, BGN, ANXA1, STAT1, CD59, CD55, DDR1, VIM, RAMP1, FGF1, THBS1 |

**Supp Table 2:** Hub genes and corresponding modules identified using Louvain clustering on the SpaceDecorr-inferred gene network in podocytes

| Module | Hubs | Genes |
| --- | --- | --- |
| 1 | VEGFA, MZT2A | MALAT1, NEAT1, MZT2A, PTGDS, ITGB8, DST, VEGFA, DCN, CD3E, CHI3L1, PSAP, TNFSF14, TYK2, SMO |
| 2 | S100A6, GSN | PLA2R1, ITGA3, ITGAV, GSN, ANXA2, SPOCK2, S100A6, HSP90B1, ST6GALNAC3, LGALS1, MXRA8, IGFBP7, BMP2, CALM2, APOD, ANXA1, VIM, ITGB5 |
| 3 | NOD2 | BATF3, CSF1R, CLEC4A, NOD2, CD28, NPR2, AHI1, CLOCK, TLR2, TUBB, YBX3, NR1H2, SMARCB1 |
| 4 | IFITM3, VIM | HLA-A, HLA-C |

**Supp Table 3:**
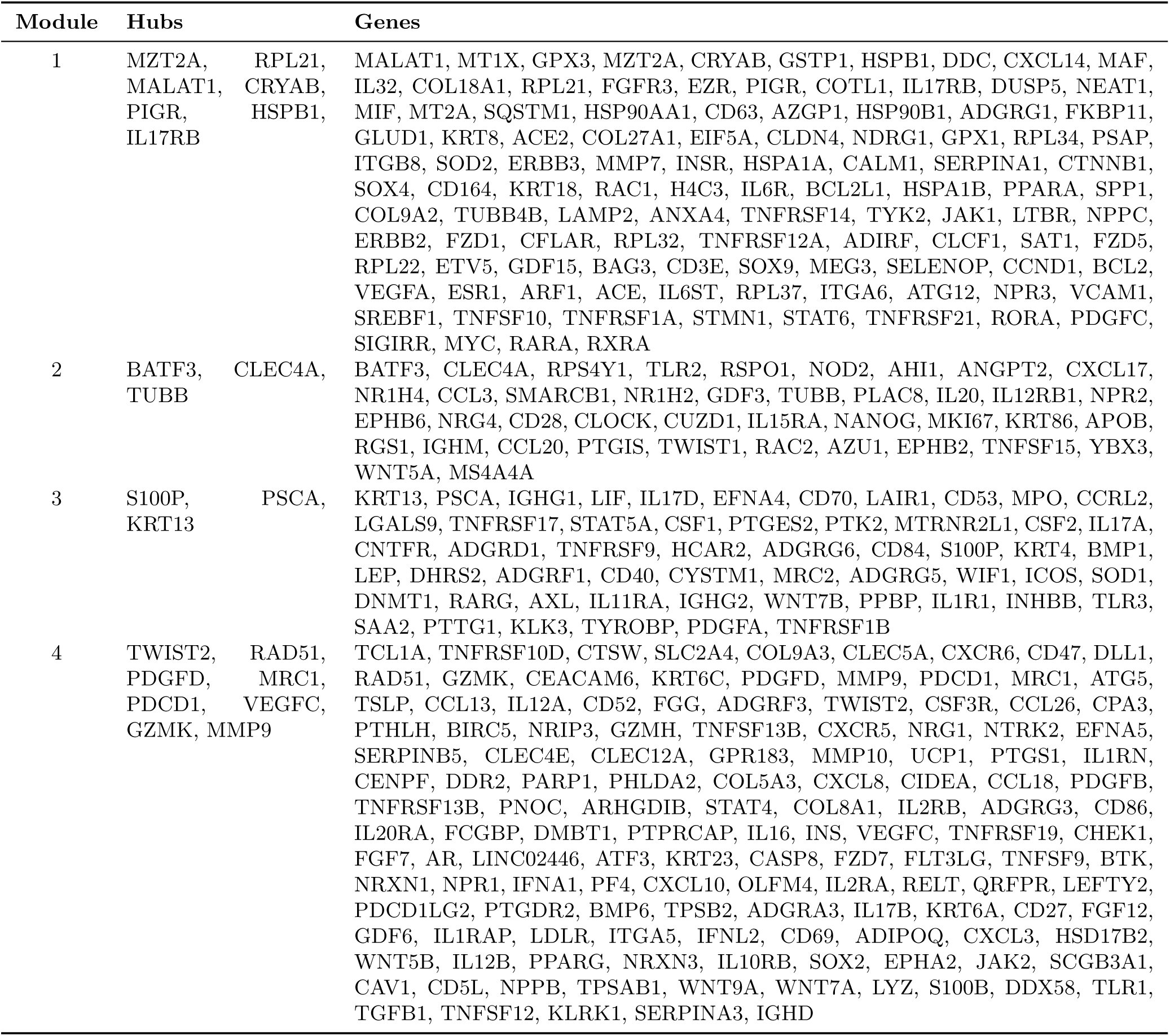
Hub genes and corresponding modules identified using Louvain clustering on the SPARK-inferred gene network in PCT

**Supp Table 4:** Hub genes and corresponding modules identified using Louvain clustering on the SpaceDecorr-inferred gene network in PCT

| Module | Hubs | Genes |
| --- | --- | --- |
| 1 | MZT2A, GPX3, CRYAB, TYK2 | MALAT1, GPX3, MT1X, MZT2A, CRYAB, MAF, DDC, CXCL14, MT2A, COL9A2, PSAP, FGFR3, PIGR, IL17RB, RPL21, NPPC, EZR, MIF, HSPB1, DUSP5, NEAT1, COL18A1, NDRG1, EIF5A, AZGP1, FKBP11, HSPA1B, CALM1, IL32, KRT8, HSPA1A, COTL1, ESR1, AZU1, SPP1, ERBB3, CD63, SERPINA1, GSTP1, COL27A1, TYK2, SQSTM1, VEGFA, TNFRSF12A, TNFRSF14, S100A9, SOD2, RPL34 |
| 2 | CLEC4A, BATF3 | BATF3, CLEC4A, TLR2, NR1H4, RSPO1, NOD2, CXCL17, RPS4Y1, ANGPT2, AHI1, TUBB, NR1H2, SMARCB1, IL20, CCL3, GDF3, NPR2, PLAC8, EPHB6, IL12RB1, CLOCK, IL15RA, CD28, NRG4, CUZD1, KRT86, APOB, MKI67, CCL20, NANOG, IGHM, TWIST1, YBX3, WNT5A, TNFSF15, PTGIS, RAC2 |
| 3 | TNFRSF17, KRT13, LIF | KRT13, LGALS9, CCRL2, PSCA, LIF, CD53, LAIR1, CD70, EFNA4, STAT5A, MPO, PTGES2, TNFRSF17, CSF1, IL17D, PTK2, IL17A, IGHG1, MTRNR2L1, TNFRSF9, CYSTM1, ADGRG5, WNT7B, WIF1, PTHLH |
| 4 | S100P | ADGRG6, HCAR2, TCL1A, ADGRF1, S100P, TNFRSF10D, CD84, CD47, LEP, CTSW, COL9A3, MRC2, DHRS2, SLC2A4, CXCR6, NRG1, IFIH1, TWIST2, IL1RN |
| 5 | TSLP, UCP1 | PDGFD, TSLP, UCP1, WNT9A |
| 6 | ADGRD1 | ADGRD1, ICOS |
| 7 | BMP1, CLEC5A | BMP1, CLEC5A |

## Notes

### Competing Interest Statement

The authors have declared no competing interest.

### Summary of Updates

Manuscript initially submitted was missing references, which were now included.

